# Tertiary and quaternary structure remodeling by occupancy of the substrate binding pocket in a large glutamate dehydrogenase

**DOI:** 10.1101/2024.12.26.630402

**Authors:** Melisa Lázaro, Nicolás Chamorro, Jorge P. López-Alonso, Diego Charro, Rodolfo M. Rasia, Gonzalo Jiménez-Osés, Mikel Valle, María-Natalia Lisa

## Abstract

Glutamate dehydrogenases (GDHs) catalyze the oxidative deamination of L-glutamate to 2-oxoglutarate using NAD(P)^+^ as a cofactor. The large type of GDHs (L-GDHs) displays a dynamic homotetrameric architecture that alternates between open and closed states. However, the catalytic mechanism and the functional relevance of the large conformational changes in L-GDHs remain poorly understood. Here, we use cryo-EM to investigate the structure and the conformational landscape of the mycobacterial L-GDH composed of 180 kDa subunits (mL-GDH_180_) when incubated with L-glutamate and NAD^+^. Classification of the heterogeneous population of tetramers reveals opening-closing motions and sorting of individual subunits resolves the occupancy of the cofactor and substrate binding pockets. Cryo-EM maps show that ligand binding to the glutamate binding pocket is accompanied by structural changes in a region approximately two nanometers away from the active site, leading to the formation of a previously undetected interaction between the catalytic domains of neighboring subunits in mL-GDH_180_ closed tetrameric states. Our findings indicate that the occupancy of the substrate binding site of mL-GDH_180_ is linked to a remodeling of both the tertiary and quaternary structure of the enzyme.

**STATEMENT FOR A BROADER AUDIENCE:** This work reveals how the binding of L-glutamate and NAD^+^ reshapes the architecture of a large glutamate dehydrogenase, linking active site occupancy to long-range structural remodeling. By capturing previously unseen conformational transitions with cryo-electron microscopy, we provide insights into the molecular logic of enzyme function in mycobacteria. These findings establish a framework to understand how structural plasticity supports metabolic control.

## INTRODUCTION

Amino acid dehydrogenases have long attracted interest due to their physiological functions and potential industrial applications. Among them, glutamate dehydrogenases (GDHs; EC 1.4.1.2) catalyze the reaction L-glutamate + NAD(P)^+^ + H_2_O <=> 2-oxoglutarate + NAD(P)H + H^+^ + NH_3_, at the crossroads of carbon and nitrogen metabolism. These enzymes are oligomeric and are classified based on the molecular weight of their subunits. Small GDHs (S-GDHs), present in prokaryotes and eukaryotes, consist of 50 kDa subunits ^1^. The conserved catalytic domain spans nearly the full length of the polypeptide chain of S-GDHs and mediates oligomerization. Biochemical and structural studies on S-GDHs have enabled detailed characterization of their catalytic and regulatory mechanisms. In contrast, much less is known about large GDHs (L-GDHs), which are found in prokaryotes and lower eukaryotes ^2^. This group includes enzymes with 115 kDa subunits (L-GDHs_115_) and those with 180 kDa subunits (L-GDHs_180_), such as the L-GDH_180_ from mycobacteria (mL-GDH_180_). L-GDHs feature distinctive and extended N- and C-terminal regions flanking the catalytic core, which may confer regulatory or structural roles ^3^.

In a previous work, we demonstrated that the L-GDH_180_ from *Mycobacterium smegmatis* (mL-GDH_180_; 76% sequence identity with the L-GDH_180_ from *Mycobacterium tuberculosis*) assembles into a homotetramer, with a quaternary structure markedly different from that of S-GDHs ^3^. Such architecture is stabilized by interactions involving the N-terminal region, which contains ACT- and PAS-type domains typically associated with regulatory functions ^4,5^, as well as a C-terminal domain of unknown function. Despite such advances, the conformational landscape explored by L-GDHs during catalysis remained poorly unknown.

Here we report cryo-EM structures of mL-GDH_180_ at resolutions up to 3 Å, obtained in the presence of NAD^+^ and L-glutamate. We reveal that ligand binding to the glutamate binding pocket is accompanied by changes in the tertiary structure in a region approximately 2 nm distant from the active site, promoting interactions between the catalytic domains of neighboring monomers and modulating the quaternary structure.

Despite extensive efforts to understand the molecular basis of GDHs activity and regulation, structural data on ternary complexes containing enzyme, cofactor, and substrate are scarce, particularly for NAD^+^-dependent L-GDHs. Here, we present cryo-EM evidence for enzyme species present upon incubation with L-glutamate and NAD^+^ of a key L-GDH_180_ for the virulence of a long-attacked pathogen ^6^, offering new insights into the conformational plasticity of L-GDHs and setting the stage for a deeper understanding of their regulation.

## RESULTS

### Three conformers of mL-GDH_180_ detected upon incubation with L-glutamate and NAD^+^ reveal modulations of the enzyme quaternary structure

To investigate the molecular basis of mL-GDH_180_ activity, we first determined its kinetic parameters for the oxidative deamination of L-glutamate. Under steady state conditions, the reaction displayed a *k*_cat_ of (15 ± 2) s^-1^; the estimated *K*_M_ values were (40 ± 10) mM for L-glutamate and (0.19 ± 0.06) mM for NAD^+^, yielding catalytic efficiencies (*k*_cat_/*K*_M_) of (0.4 ± 0.1) mM^-1^ s^-1^ and (80 ± 30) mM^-1^ s^-1^, respectively (Figure S1A), as determined in 50 mM Hepes, 50 mM NaCl, at pH 6.8, near the apparent pH optimum of the reaction (Figure S1B). Single-particle cryo-EM experiments were then conducted on mL-GDH_180_ incubated with NAD^+^ and L-glutamate under NADH-producing conditions, during early turnover (Figure S1C; see also Materials and methods). As previously reported ^3^, 2D class averages revealed high flexibility at the distal ends of the tetramers, where the ACT*1-2 and PAS domains are located (Figure S2). This flexibility resulted in vanishing density for residues 1-499 in the cryo-EM maps. Subsequently, 3D-classification of mL-GDH_180_ particles revealed three conformational states of the enzyme, named Open (32.4%), Closed1 (39.6%) and Closed2 (28%) (Figure 1, Table I, Figure S3). In these conformers, the ACT*3 module, the helical motif 3 (HM3), the catalytic domain and the C-terminal region were defined in each monomer, achieving estimated global resolutions of 3.88 Å, 3.54 Å and 3.57 Å, respectively. The conformers differ primarily in the relative positions of the subunits (Figure 1A, Figure S4A and Movie S1). In the Closed forms, the catalytic domains of adjacent subunits connected through their N-terminal segments are found in closer proximity compared to the Open state (Figure 1B).

**Figure 1.**
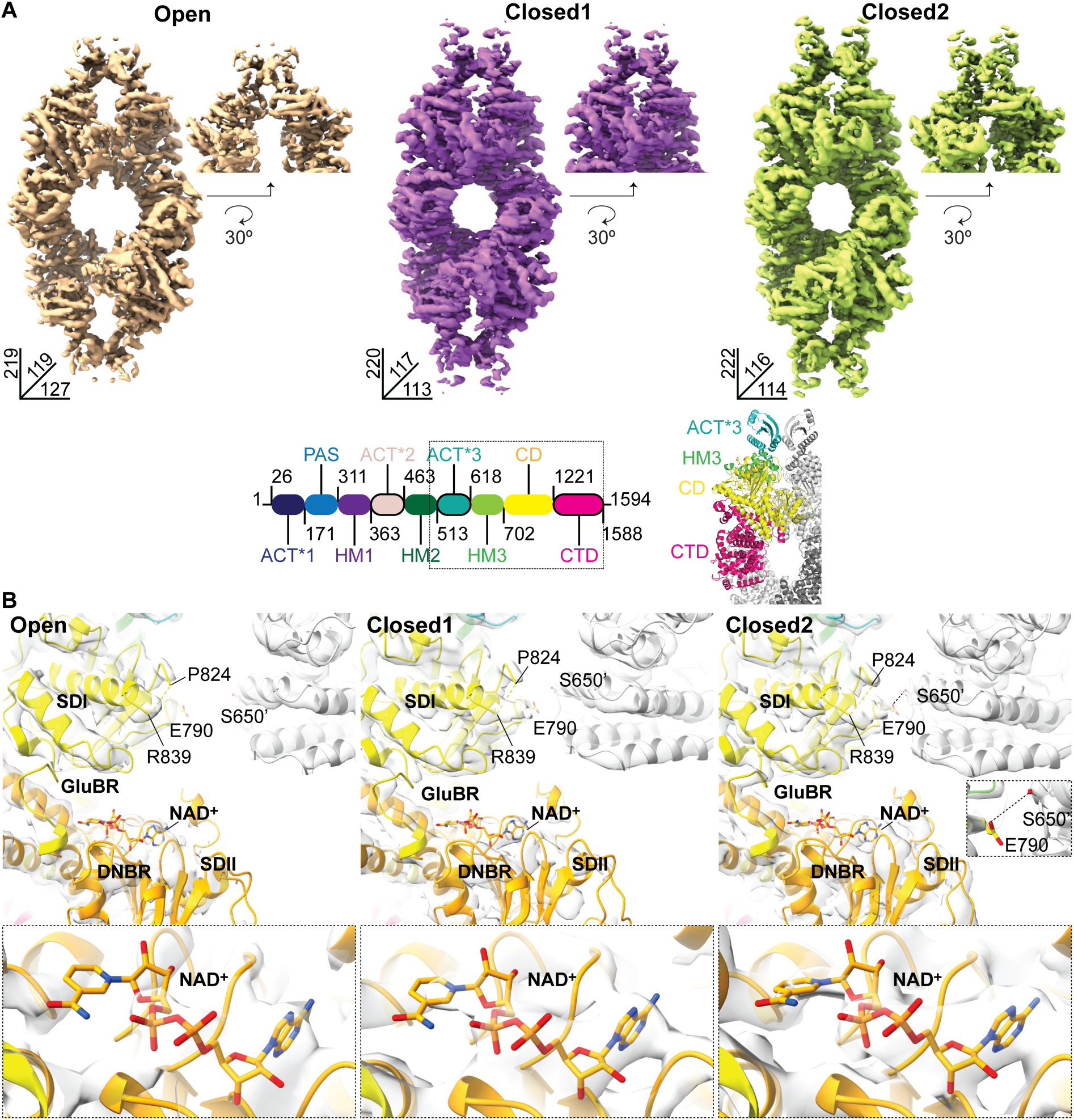
Three conformers of mL-GDH_180_ detected upon incubation with L-glutamate and NAD^+^ reveal modulations of the enzyme quaternary structure. (A) Cryo-EM mL-GDH_180_ maps obtained after incubating the enzyme with L-glutamate and NAD^+^. The dimensions of each conformer are provided in Å. Maps cover, for each monomer, the region indicated by a dotted rectangle in the diagram shown below (residues 500-1588). ACT*, Aspartate kinase-Chorismate mutase-TyrA-like; PAS, Per-Arnt-Sim; HM, helical motif; CD, catalytic domain; CTD, C-terminal domain. (B) Approach of the catalytic domains of mL-GDH_180_ subunits linked by the N-terminal region. Segment 825-838, poorly defined in maps of the Open, Closed1 and Closed2 enzyme forms, approaches the catalytic domain in a neighboring subunit. In each case, a zoomed in view of the dinucleotide binding region (DNBR) is shown below. The protein is displayed in ribbon representation and cryo-EM maps are depicted in grey. The nicotinamide portion of the NAD^+^ molecule (in sticks) exhibits high mobility in all conformations. Subdomains I and II (SDI and SDII) of the catalytic domain are colored in yellow and orange, respectively, whereas ACT*3 and HM3 motifs are shown in cyan and green, respectively, for one monomer (the one on the left), with the neighboring subunit (the one on the right) shown in grey. All maps are shown with a contour level of 0.02. GluBR, glutamate binding region.

**Table I.**
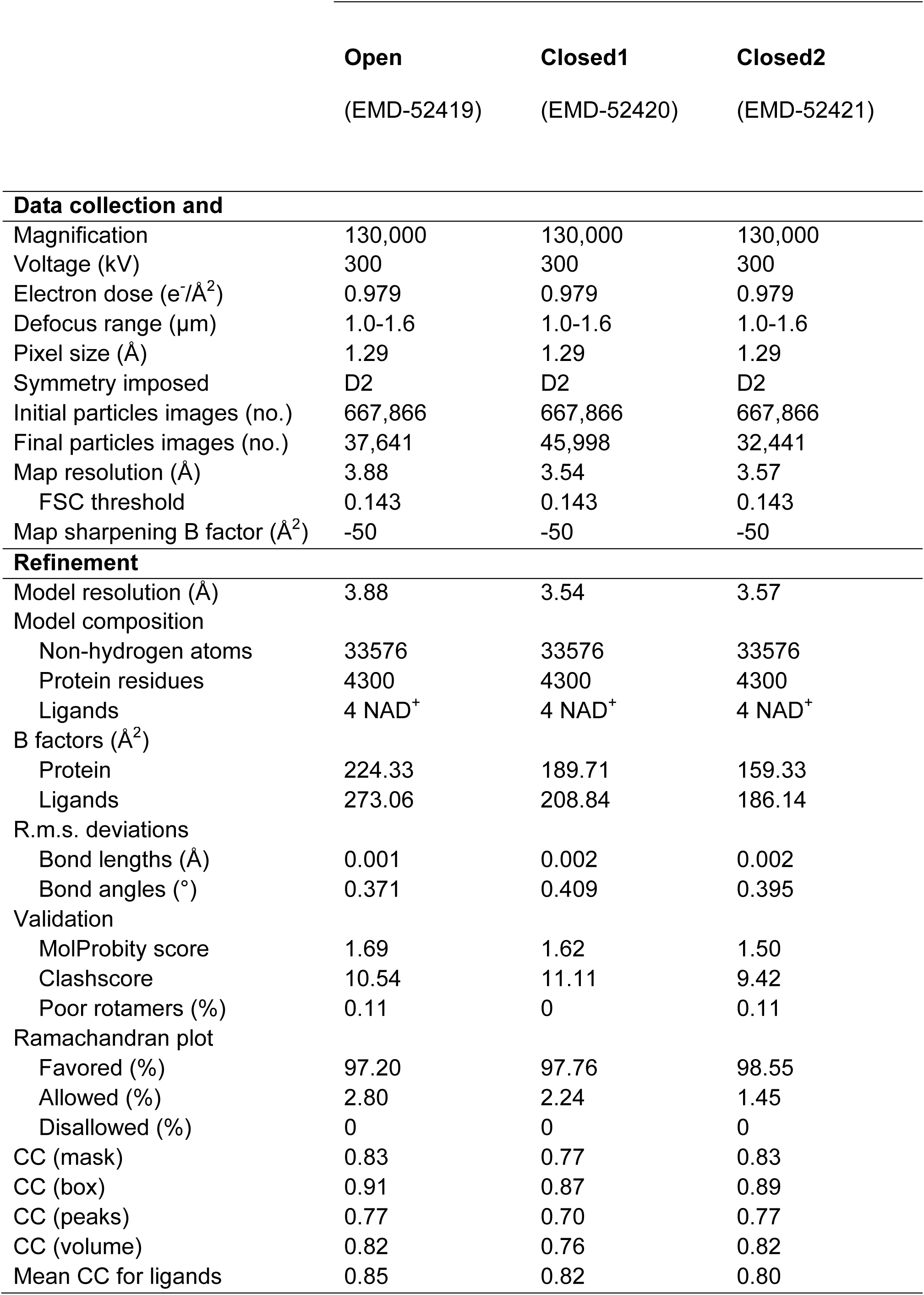
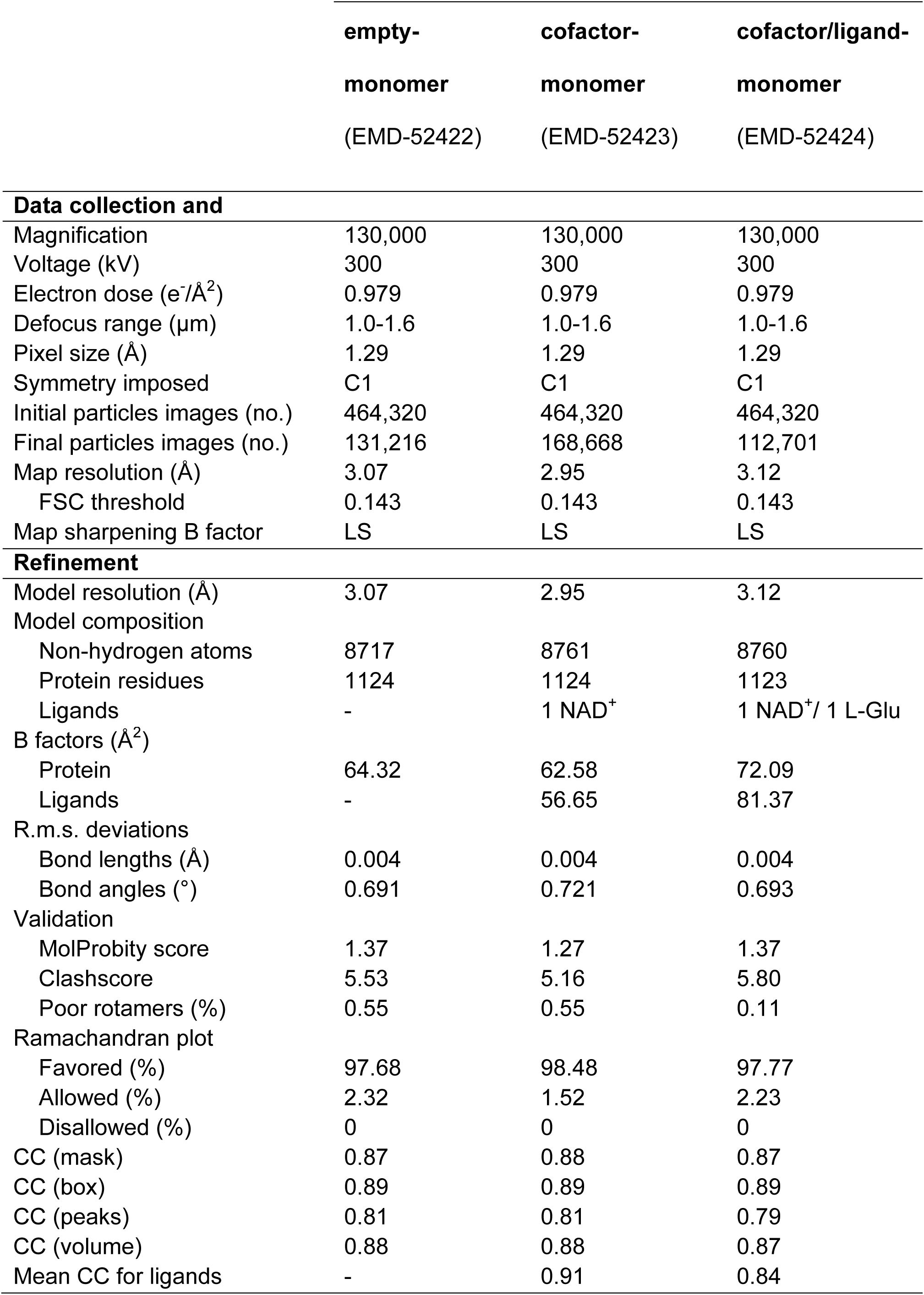
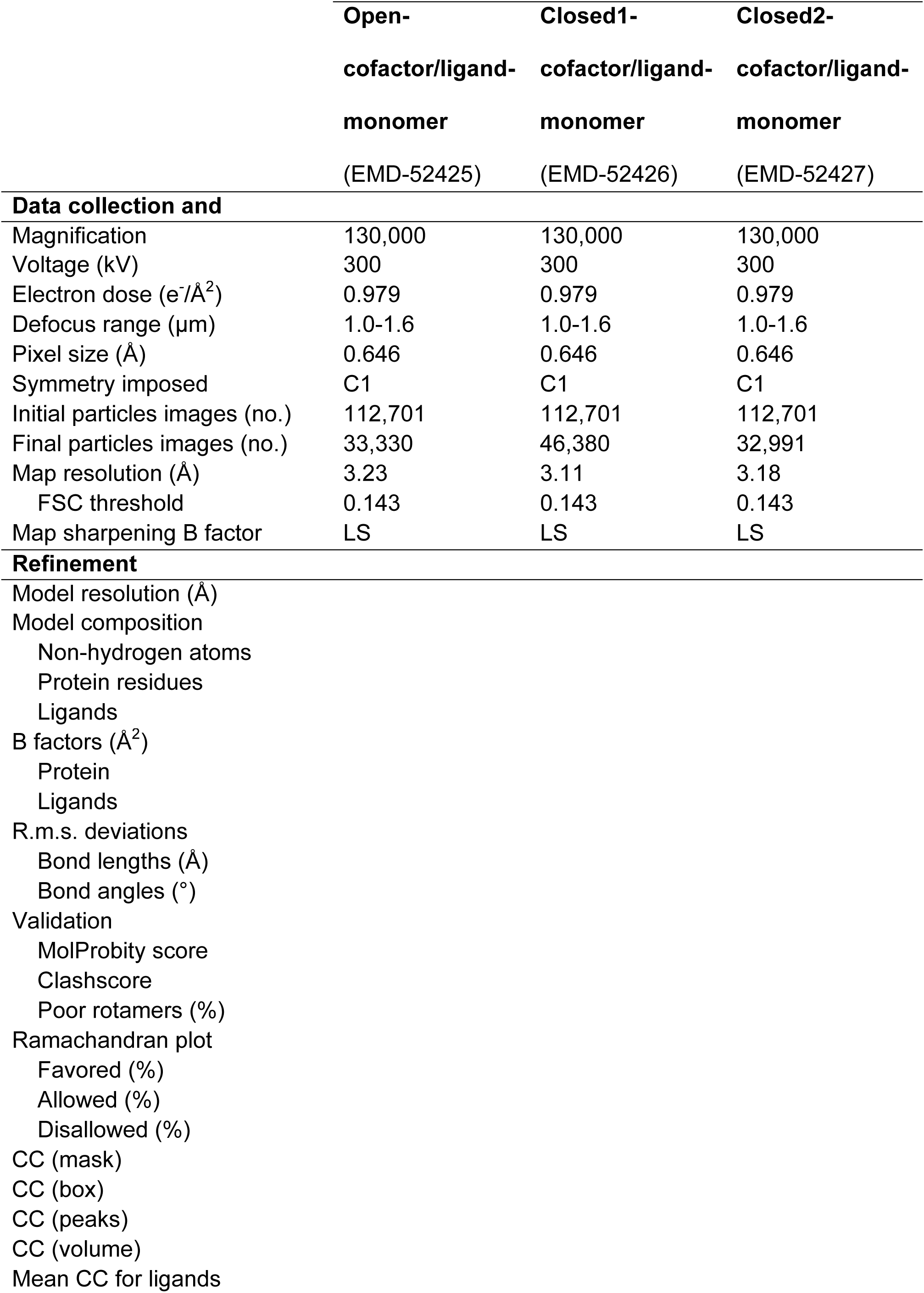
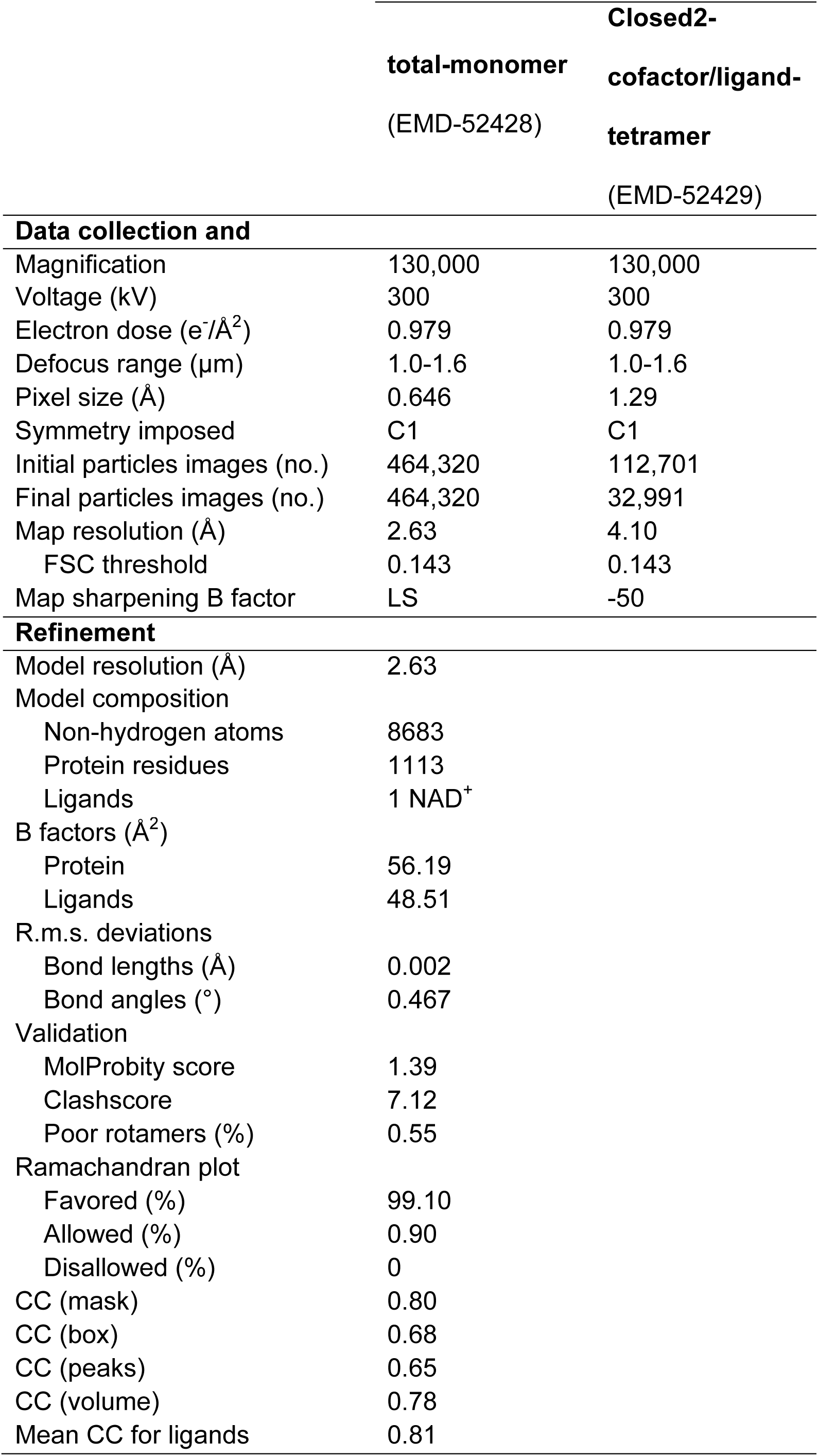
Cryo-EM data collection and processing. LS, local sharpening.

|  | **Open**<br>(EMD-52419) | **Closed1**<br>(EMD-52420) | **Closed2**<br>(EMD-52421) |
| --- | --- | --- | --- |
| **Data collection and** |  |  |  |
| Magnification | 130,000 | 130,000 | 130,000 |
| Voltage (kV) | 300 | 300 | 300 |
| Electron dose (e<sup>-</sup>/Å<sup>2</sup>) | 0.979 | 0.979 | 0.979 |
| Defocus range (μm) | 1.0-1.6 | 1.0-1.6 | 1.0-1.6 |
| Pixel size (Å) | 1.29 | 1.29 | 1.29 |
| Symmetry imposed | D2 | D2 | D2 |
| Initial particles images (no.) | 667,866 | 667,866 | 667,866 |
| Final particles images (no.) | 37,641 | 45,998 | 32,441 |
| Map resolution (Å) | 3.88 | 3.54 | 3.57 |
| FSC threshold | 0.143 | 0.143 | 0.143 |
| Map sharpening B factor (Å<sup>2</sup>) | -50 | -50 | -50 |
| **Refinement** |  |  |  |
| Model resolution (Å) | 3.88 | 3.54 | 3.57 |
| Model composition |  |  |  |
| Non-hydrogen atoms | 33576 | 33576 | 33576 |
| Protein residues | 4300 | 4300 | 4300 |
| Ligands | 4 NAD<sup>+</sup> | 4 NAD<sup>+</sup> | 4 NAD<sup>+</sup> |
| B factors (Å<sup>2</sup>) |  |  |  |
| Protein | 224.33 | 189.71 | 159.33 |
| Ligands | 273.06 | 208.84 | 186.14 |
| R.m.s. deviations |  |  |  |
| Bond lengths (Å) | 0.001 | 0.002 | 0.002 |
| Bond angles (°) | 0.371 | 0.409 | 0.395 |
| Validation |  |  |  |
| MolProbity score | 1.69 | 1.62 | 1.50 |
| Clashscore | 10.54 | 11.11 | 9.42 |
| Poor rotamers (%) | 0.11 | 0 | 0.11 |
| Ramachandran plot |  |  |  |
| Favored (%) | 97.20 | 97.76 | 98.55 |
| Allowed (%) | 2.80 | 2.24 | 1.45 |
| Disallowed (%) | 0 | 0 | 0 |
| CC (mask) | 0.83 | 0.77 | 0.83 |
| CC (box) | 0.91 | 0.87 | 0.89 |
| CC (peaks) | 0.77 | 0.70 | 0.77 |
| CC (volume) | 0.82 | 0.76 | 0.82 |
| Mean CC for ligands | 0.85 | 0.82 | 0.80 |

|  | **empty-**<br>**monomer**<br>(EMD-52422) | **cofactor-**<br>**monomer**<br>(EMD-52423) | **cofactor/ligand-**<br>**monomer**<br>(EMD-52424) |
| --- | --- | --- | --- |
| **Data collection and** |  |  |  |
| Magnification | 130,000 | 130,000 | 130,000 |
| Voltage (kV) | 300 | 300 | 300 |
| Electron dose (e<sup>-</sup>/Å<sup>2</sup>) | 0.979 | 0.979 | 0.979 |
| Defocus range (μm) | 1.0-1.6 | 1.0-1.6 | 1.0-1.6 |
| Pixel size (Å) | 1.29 | 1.29 | 1.29 |
| Symmetry imposed | C1 | C1 | C1 |
| Initial particles images (no.) | 464,320 | 464,320 | 464,320 |
| Final particles images (no.) | 131,216 | 168,668 | 112,701 |
| Map resolution (Å) | 3.07 | 2.95 | 3.12 |
| FSC threshold | 0.143 | 0.143 | 0.143 |
| Map sharpening B factor | LS | LS | LS |
| **Refinement** |  |  |  |
| Model resolution (Å) | 3.07 | 2.95 | 3.12 |
| Model composition |  |  |  |
| Non-hydrogen atoms | 8717 | 8761 | 8760 |
| Protein residues | 1124 | 1124 | 1123 |
| Ligands | - | 1 NAD<sup>+</sup> | 1 NAD<sup>+</sup>/ 1 L-Glu |
| B factors (Å<sup>2</sup>) |  |  |  |
| Protein | 64.32 | 62.58 | 72.09 |
| Ligands | - | 56.65 | 81.37 |
| R.m.s. deviations |  |  |  |
| Bond lengths (Å) | 0.004 | 0.004 | 0.004 |
| Bond angles (°) | 0.691 | 0.721 | 0.693 |
| Validation |  |  |  |
| MolProbity score | 1.37 | 1.27 | 1.37 |
| Clashscore | 5.53 | 5.16 | 5.80 |
| Poor rotamers (%) | 0.55 | 0.55 | 0.11 |
| Ramachandran plot |  |  |  |
| Favored (%) | 97.68 | 98.48 | 97.77 |
| Allowed (%) | 2.32 | 1.52 | 2.23 |
| Disallowed (%) | 0 | 0 | 0 |
| CC (mask) | 0.87 | 0.88 | 0.87 |
| CC (box) | 0.89 | 0.89 | 0.89 |
| CC (peaks) | 0.81 | 0.81 | 0.79 |
| CC (volume) | 0.88 | 0.88 | 0.87 |
| Mean CC for ligands | - | 0.91 | 0.84 |

|  | **Open-** | **Closed1-** | **Closed2-** |
| --- | --- | --- | --- |
|  | **cofactor/ligand-** | **cofactor/ligand-** | **cofactor/ligand-** |
|  | **monomer** | **monomer** | **monomer** |
|  | (EMD-52425) | (EMD-52426) | (EMD-52427) |
| **Data collection and** |  |  |  |
| Magnification | 130,000 | 130,000 | 130,000 |
| Voltage (kV) | 300 | 300 | 300 |
| Electron dose (e<sup>-</sup>/Å<sup>2</sup>) | 0.979 | 0.979 | 0.979 |
| Defocus range (μm) | 1.0-1.6 | 1.0-1.6 | 1.0-1.6 |
| Pixel size (Å) | 0.646 | 0.646 | 0.646 |
| Symmetry imposed | C1 | C1 | C1 |
| Initial particles images (no.) | 112,701 | 112,701 | 112,701 |
| Final particles images (no.) | 33,330 | 46,380 | 32,991 |
| Map resolution (Å) | 3.23 | 3.11 | 3.18 |
| FSC threshold | 0.143 | 0.143 | 0.143 |
| Map sharpening B factor | LS | LS | LS |
| **Refinement** |  |  |  |
| Model resolution (Å) |  |  |  |
| Model composition |  |  |  |
| Non-hydrogen atoms |  |  |  |
| Protein residues |  |  |  |
| Ligands |  |  |  |
| B factors (Å<sup>2</sup>) |  |  |  |
| Protein |  |  |  |
| Ligands |  |  |  |
| R.m.s. deviations |  |  |  |
| Bond lengths (Å) |  |  |  |
| Bond angles (°) |  |  |  |
| Validation |  |  |  |
| MolProbity score |  |  |  |
| Clashscore |  |  |  |
| Poor rotamers (%) |  |  |  |
| Ramachandran plot |  |  |  |
| Favored (%) |  |  |  |
| Allowed (%) |  |  |  |
| Disallowed (%) |  |  |  |
| CC (mask) |  |  |  |
| CC (box) |  |  |  |
| CC (peaks) |  |  |  |
| CC (volume) |  |  |  |
| Mean CC for ligands |  |  |  |

|  | **total-monomer**<br>(EMD-52428) | **Closed2-cofactor/ligand-tetramer**<br>(EMD-52429) |
| --- | --- | --- |
| **Data collection and** |  |  |
| Magnification | 130,000 | 130,000 |
| Voltage (kV) | 300 | 300 |
| Electron dose (e<sup>-</sup>/Å<sup>2</sup>) | 0.979 | 0.979 |
| Defocus range (μm) | 1.0-1.6 | 1.0-1.6 |
| Pixel size (Å) | 0.646 | 1.29 |
| Symmetry imposed | C1 | C1 |
| Initial particles images (no.) | 464,320 | 112,701 |
| Final particles images (no.) | 464,320 | 32,991 |
| Map resolution (Å) | 2.63 | 4.10 |
| FSC threshold | 0.143 | 0.143 |
| Map sharpening B factor | LS | -50 |
| **Refinement** |  |  |
| Model resolution (Å) | 2.63 |  |
| Model composition |  |  |
| Non-hydrogen atoms | 8683 |  |
| Protein residues | 1113 |  |
| Ligands | 1 NAD<sup>+</sup> |  |
| B factors (Å<sup>2</sup>) |  |  |
| Protein | 56.19 |  |
| Ligands | 48.51 |  |
| R.m.s. deviations |  |  |
| Bond lengths (Å) | 0.002 |  |
| Bond angles (°) | 0.467 |  |
| Validation |  |  |
| MolProbity score | 1.39 |  |
| Clashscore | 7.12 |  |
| Poor rotamers (%) | 0.55 |  |
| Ramachandran plot |  |  |
| Favored (%) | 99.10 |  |
| Allowed (%) | 0.90 |  |
| Disallowed (%) | 0 |  |
| CC (mask) | 0.80 |  |
| CC (box) | 0.68 |  |
| CC (peaks) | 0.65 |  |
| CC (volume) | 0.78 |  |
| Mean CC for ligands | 0.81 |  |

Comparison of the atomic models derived from the 3D cryo-EM maps provided initial insights on the molecular determinants underlying the modulation of mL-GDH_180_ architecture. Pairwise structural alignments of monomers from the three conformers, based on all modeled α-carbons, yielded an average RMSD (root-mean-square deviation) of 0.4 Å (calculated over 1075 aligned residues), indicating that the observed conformational variability primarily reflects quaternary rearrangements.

Consistent with this, the intersubunit contact area at the AB-CD interface (Figure S4B) was measured ^7^ as 550 Å^2^, 750 Å^2^ and 800 Å^2^, and at the AD-BC interface as 1200 Å^2^, 1000 Å^2^ and 1200 Å^2^, for the Open, Closed1 and Closed2 forms, respectively. At the AB-CD interface, hydrogen bonds between residues in the ACT*3 domains of neighboring monomers mediate the intersubunit contacts in all three conformers (Figure S4B and Table SI). Remarkably, a previously undetected interfacial interaction was found in the Closed2 tetramer, involving a hydrogen bond between the side chains of Ser650 (in the HM3) and Glu790 (in the catalytic domain) (Figure 1B and Figure S4B). At the AD-BC interface, distinct salt bridge combinations are found in the different conformers, with Asp1516:Arg1582 present in the Open form, Arg1504:Glu1562 in the Closed1 and both simultaneously observed in the Closed2 (Figure S4B and Table SI). Additionally, in all three conformations, extensive hydrogen bonds networks link a conserved set of secondary structure elements in the C-terminal domain, with alternative arrangements in each state. Most of the interfacial residues observed in mL-GDH_180_ tetramers found in the presence of NAD^+^ and L-glutamate had been previously identified as contributors to oligomer stabilization ^3^ and are conserved across diverse L-GDHs (Figure S5), suggesting a shared structural basis for quaternary modulation within this enzyme family.

As HM3 in one monomer and the region encompassing Glu790 in the catalytic domain of an adjacent subunit move closer, other portions of the catalytic domains in mL-GDH_180_ monomers linked via their N-terminal segments also shift toward each other (Figure 1B). Intriguingly, the density corresponding to residues 825-838 was poorly defined in the maps of the Open, Closed1 and Closed2 conformers distinguished in the presence of NAD^+^ and L-glutamate. At the same time, the nicotinamide moiety of the cofactor molecule placed within the active site appeared highly mobile in all three species, although this mobility seemed reduced in the Closed1 and Closed2 forms. No additional density was detected within the active sites beyond that attributable to the cofactor.

### Three conformers of mL-GDH_180_ subunits reveal modulations in tertiary structure associated with differential active site occupancies

To clarify the absence of signal for any ligand other than the cofactor in the cryo-EM maps of mL-GDH_180_ tetramers, we performed a hierarchical classification of all mL-GDH_180_ subunits, regardless of whether they belonged to the Open, Closed1 or Closed2 conformers, using a mask that covered the catalytic domain (Figure S3).

This classification yielded three distinct forms for mL-GDH_180_ monomers, termed empty-monomer, cofactor-monomer and cofactor/ligand-monomer, with estimated resolutions of 3.07 Å, 2.95 Å and 3.12 Å, respectively (Figure 2 and Table I). The empty-monomer map shows faint density within the cofactor binding pocket, but does not reveal strong binding to that site, nor any signal in the substrate binding pocket.

**Figure 2.**
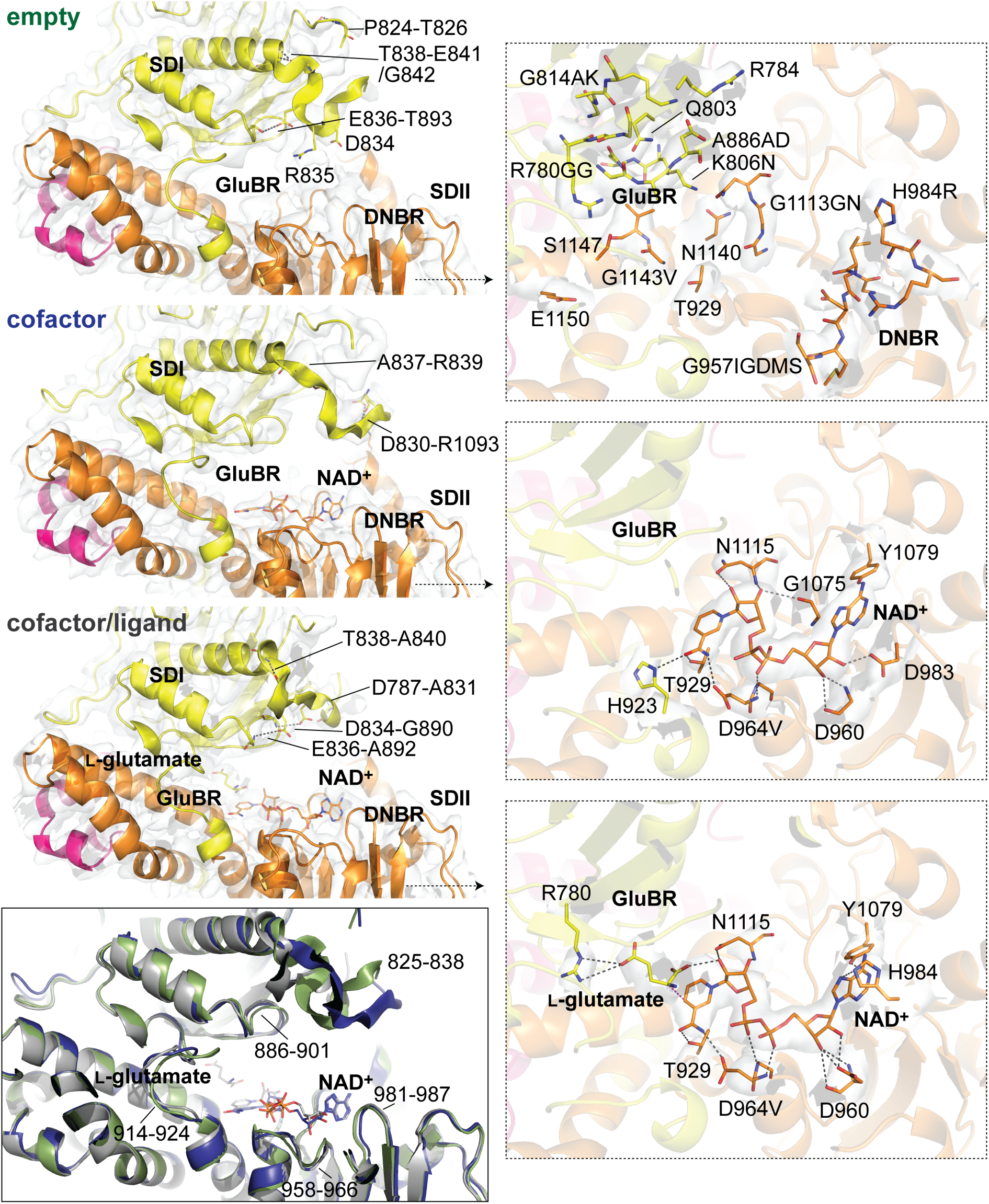
Three conformers of mL-GDH_180_ subunits reveal modulations in tertiary structure associated with differential active site occupancies. Alternative conformations of mL-GDH_180_ subunits are shown, with the active site either empty or containing NAD^+^ or NAD^+^ and L-glutamate. The average RMSD for 1122 aligned residues in pairwise comparisons of the atomic models (show in ribbon representation) is 0.7 Å. NAD^+^ and L-glutamate are depicted as sticks as well as highlighted residues. For each species, subdomains I and II of the catalytic domain (SDI, residues 702-926 and 1188-1220; SDII, residues 927-1187) are colored in yellow and orange, respectively. A small portion of the C-terminal domain is displayed in magenta. Interactions established by the segment 825-838 with other portions of the polypeptide chain are shown in the panels on the left. Note that within the region 825–838, residues 825-827 and 830-838 could be built for the empty- and cofactor-monomers, while residues 825 and 829-838 were modeled for the cofactor/ligand-monomer. A comparison of the atomic models of the empty- (green), cofactor- (blue) and cofactor/ligand-monomer (gray) is presented in the lower left corner. Cryo-EM maps are depicted in grey, with contour levels of 5. H-bonds and salt bridges are displayed as dashed lines. GluBR, glutamate binding region; DNBR, dinucleotide binding region.

In contrast, the cofactor-monomer map clearly displays a bound cofactor molecule in the catalytic cavity and the cofactor/ligand-monomer map discloses both the cofactor and an additional density element within the active site (Figure 2 and Figure S6). An average of all these variants resulted in a 2.63 Å resolution map of the mL-GDH_180_ monomer (Figure S6 and Figure S7), which was used to model the polypeptide segment comprising residues 463–1588 before refining the atomic coordinates for each occupancy state.

The atomic model derived from the empty-monomer map revealed the arrangement of residues that, according to previous analyses ^2,3^, define the dinucleotide binding region (DNBR) and the glutamate binding region (GluBR) of mL-GDH_180_ (Figure 2). The cryo-EM maps of the cofactor- and cofactor/ligand-monomers of mL-GDH_180_ allowed to place a NAD^+^ molecule in extended conformation within the DNBR (Figure 2). According to our models, the pyrophosphate moiety is located in the crevice formed between strands β1 and β5 (residues 953-958 and 1070-1073, respectively) of subdomain II of the catalytic domain (SDII) and is stabilized by the dipole of the helix comprising residues 964-969, which connects strands β1 and β2 (residues 976-982) and is known as the dinucleotide binding helix (Figure 3). The adenine ring of NAD^+^ fits well in the density, the glycosidic bond to the nicotinamide ring adopts a *syn* conformation and both riboses display C2’ *endo* puckering, in agreement with previous reports for S-GDHs ^8^.

**Figure 3.**
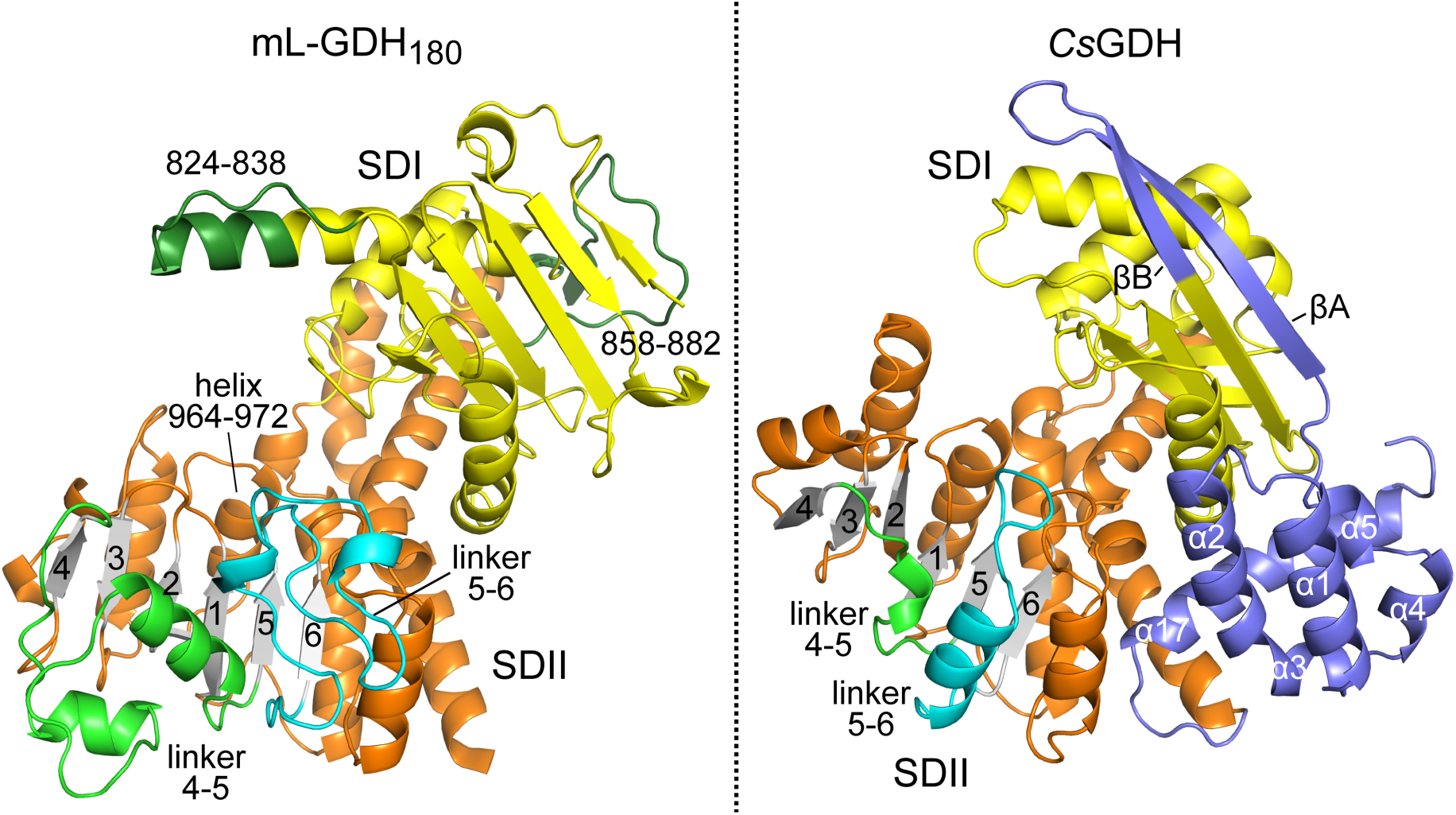
Comparison of the catalytic domain of mL-GDH_180_ with that of S-GDHs. A monomer of mL-GDH_180_ is shown on the left. Segments 824-838 and 858-882 (in dark green), both located in the catalytic domain subdomain I (SDI), are absent in S-GDHs ^2^. For the 824-838 portion coordinates were taken from a model calculated with AlphaFold 3 ^33^. A monomer of *Cs*GDH (*Cs*, *Clostridium symbiosum*; PDB 1BGV ^20^), considered a representative S-GDH, is shown on the right. The SDI of S-GDHs presents secondary structure elements (in dark blue, named according to the consensus nomenclature for S-GDHs) with conserved roles in oligomerization in the enzyme subfamily, which are not present in mL-GDH_180_ or related L-GDHs. For both mL-GDH_180_ and *Cs*GDH, the six-stranded 432156 β-sheet that shapes the DNBR in the catalytic domain subdomain II (SDII) is shown in grey; the linker regions β4-β5 (in light green) and β5-β6 (in cyan) encompass a larger number of residues in mL-GDH_180_. For the two proteins, the SDI and SDII are shown, except for otherwise highlighted portions, in yellow and orange, respectively.

Notably, the pattern of NAD^+^-protein interactions differs between cofactor- and cofactor/ligand-mL-GDH_180_ monomers (Figure 2). In the cofactor-monomer, the adenine ring interacts with the side chain (sc) of Tyr1079, the 2’ and 3’ hydroxyl groups of the adenosine ribose point towards the surface of the protein and hydrogen bond Asp983 (sc) and the main chain (mc) of Asp960, respectively, phosphates interact with the amide nitrogen of Asp964 and Val965, the nicotinamide ribose hydrogen bonds Gly1075 (mc) and Asn1115 (mc and sc), and the nicotinamide amide contacts His923 (sc), Thr929 (sc) and Asp964 (sc). In contrast, in the cofactor/ligand-monomer, the adenine of NAD^+^ interacts with Tyr1079 (sc) as well as His984 (sc), both adenosine hydroxyls hydrogen bond Asp960 (sc and mc), the nicotinamide ribose interacts only with Asn1115 (mc and sc) and the nicotinamide amide contacts only Thr929 (sc) and Asp964 (sc). These differences, along with subtle but detectable conformational changes in segments 958-966 and 981-987 that contribute to the DNBR, correlate with a shift of the cofactor toward the GluBR in the cofactor/ligand-monomer. In this state, a novel density element becomes evident (Movie S2), within which it is possible to position a substrate molecule (CC = 0.79) such that the α-carbon of the amino acid lies 3.3 Å from the C4 atom of the nicotinamide ring of the cofactor, explaining the continuous density observed between them. Hence, the γ-carboxylate of the ligand forms salt bridges with Arg780 (sc).

Strikingly, segment 914-924, which lines the active site cleft, and region 886-901, which borders the glutamate binding pocket, adopt distinct conformations in the empty-, cofactor- and cofactor/ligand-mL-GDH_180_ monomers (Figure 2). These variations are associated with alternate conformations of segment 825-838, which is better defined in the maps of the monomers than in the tetrameric reconstructions. Thus, residues 825-827 and 830-838 could be built for the empty- and cofactor-monomers, while residues 825 and 829-838 were modeled for the cofactor/ligand-monomer. In the empty-monomer, Arg835 is directed toward Asp964 and Asp960, while Asp834 faces His984. When the cofactor binds within the DNBR, engaging residues Asp964, Asp960 and His984, Arg835 and Asp834 shift from their positions and the segment 830-838 loosens its packing against the catalytic core. When the GluBR is also occupied, segment 831-837 packs against residues 889-894. The amino acid string 825-838 of mL-GDH_180_ is found in L-GDHs_180_ but not in related low molecular weight enzymes (Figure 3 and Figure S5) ^9^.

### GluBR occupancy is linked to the coupling of the catalytic domains of two adjacent monomers in the Closed mL-GDH_180_ conformations

To elucidate structural changes in the context of mL-GDH_180_ tetramers, we next discriminated the monomeric conformers according to whether they originated from Open, Closed1 and Closed2. This analysis revealed nine distinct species: Open-empty-monomer, Closed1-empty-monomer, Closed2-empty-monomer (3.5 Å, 3.35 Å and 3.48 Å resolution, respectively), Open-cofactor-monomer, Closed1-cofactor-monomer, Closed2-cofactor-monomer (3.3 Å, 3.23 Å and 3.28 Å), Open-cofactor/ligand-monomer, Closed1-cofactor/ligand-monomer and Closed2-cofactor/ligand-monomer (3.23 Å, 3.11 Å and 3.18 Å) (Figure S3).

No discernable structural differences were observed among the empty- or cofactor-monomers across the tetrameric states. In contrast, cryo-EM maps of the cofactor/ligand-monomers displayed heterogeneity within the glutamate binding pocket. In the Open-cofactor/ligand-monomer map, a continuous density extends from the cofactor nicotinamide ring into the GluBR (Figure 4, see also Figure 2), although it fades in the region where the γ-carboxylate of the substrate would be stabilized. Conversely, the maps of the Closed1- and Closed2-cofactor/ligand monomers display ligand density within the GluBR primarily near Arg780, with a weaker signal in the Closed2 state.

**Figure 4.**
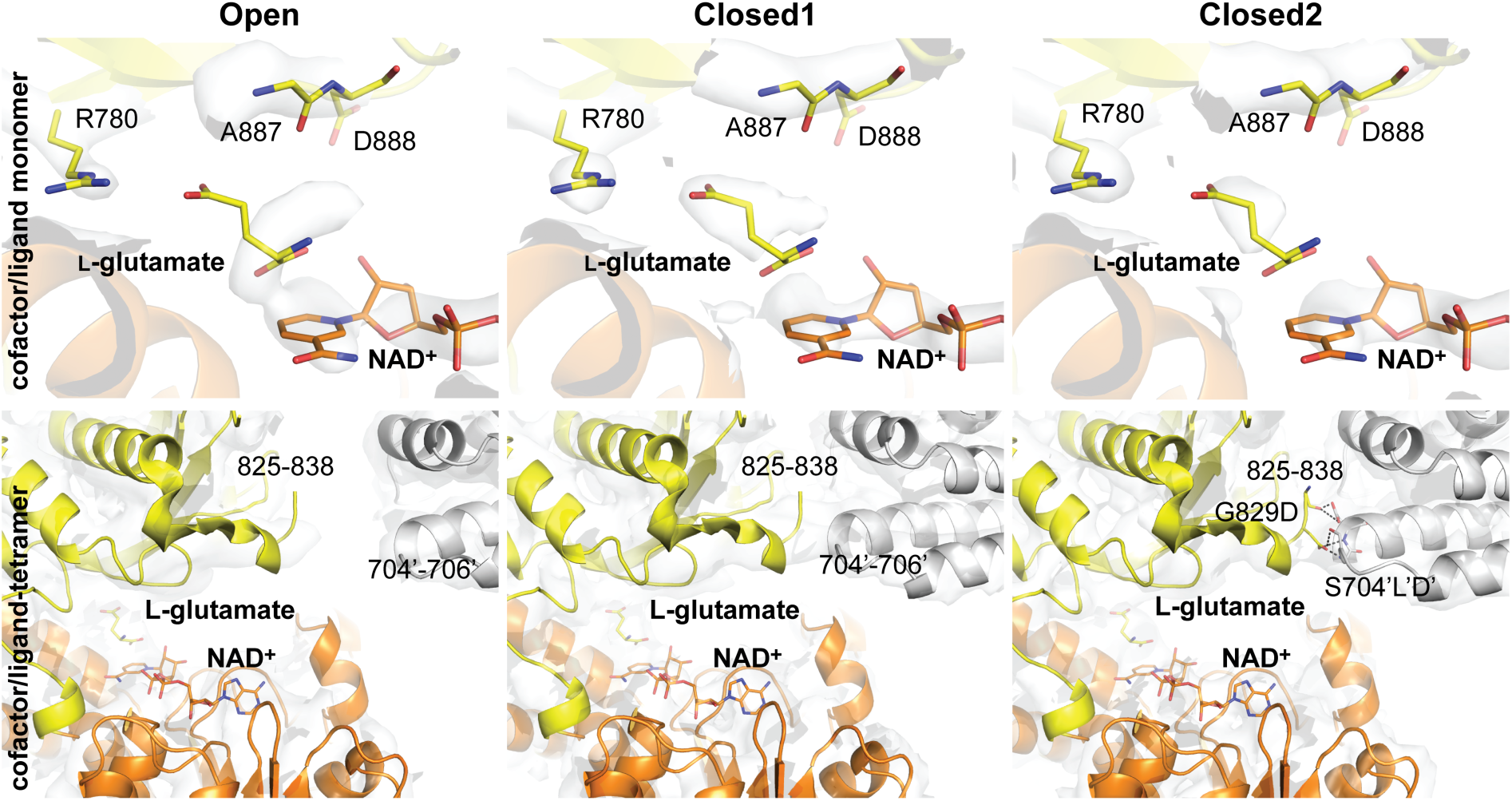
GluBR occupancy is linked to the coupling of the catalytic domain of two adjacent monomers in the Closed conformations. Top panel: the atomic model of the cofactor/ligand-monomer of mL-GDH_180_ is shown together with maps of this species obtained from Open, Closed1 or Closed2 tetramers (Open-, Closed1- and Closed2-cofactor/ligand-monomers; contour levels are 5). NAD^+^ and L-glutamate as well as highlighted residues are depicted as sticks. The subdomains I and II of the catalytic domain (SDI and SDII) are colored in yellow and orange, respectively. Bottom panel: in the context of tetramers containing a cofactor/ligand-monomer (cofactor/ligand-tetramers; contour levels are 0.015), in a progression towards the Closed1 and Closed2 conformers, the stabilization of an interaction between segment 825-838 of one subunit (colored, on the left) and segment 704-706 of the neighboring one (in grey, on the right) is observed. The atomic coordinates of the subunit on the left correspond in each case to those of a cofactor/ligand-monomer, replacing the equivalent ones presented in Figure 1. H-bonds and salt bridges are displayed as dashed lines. Cryo-EM maps are displayed in grey.

As noted earlier, incubation of mL-GDH_180_ with NAD^+^ and L-glutamate induces remodeling of the region 825-838 (Figure 2). Strikingly, this segment interacts with residues 704-706 of the catalytic domain of a neighboring N-terminally-linked subunit, giving rise to an additional molecular bridge in the Closed tetramers (Figure 4). This intersubunit interaction appears strengthened in the Closed2 state (Table I).

Importantly, it is only observed when the monomer contributing the 825-838 region has both its cofactor and substrate binding pockets simultaneously occupied. The interaction occurs approximately 2 nm from the active site of the ligand-loaded monomer (Movie S3), linking GluBR occupancy to the physical contact between adjacent catalytic domains and suggesting a structural basis for turnover regulation.

## DISCUSSION

GDHs constitute a large family of enzymes that function in all kingdoms of life at the crossroad between the Krebs cycle and ammonium assimilation. While S-GDHs have been studied for over half a century, it was not until 2021 that the first experimental evidence of the 3D architecture of an L-GDH was obtained ^3^. In that work, we determined that the quaternary structure of mL-GDH_180_ differs radically from that of S-GDHs, and that such assembly is shaped by the distinctive N- and C-terminal extensions characteristic of L-GDHs ^3^. Here we present cryo-EM data reaching resolutions up to 2.63 Å for mL-GDH_180_ (Table I), captured in the presence of NAD^+^ during the oxidative deamination of L-glutamate.

Our 3D classification of subunits successfully separated mL-GDH_180_ monomers with both the DNBR and GluBR simultaneously occupied, as well as monomers bound to NAD^+^ alone (Figure 1, Figure 2 and Figure 4). The NAD^+^ binding mode in mL-GDH_180_ (Figure 2 and Figure 4) mirrors that described for *Cs*GDH (S-GDH from *Clostridium symbiosum*) ^8^. The adenine ring is stabilized by hydrogen bonds with His984 (Pro262 in *Cs*GDH) and Tyr1079 (Asp325), the NH and CO groups of the nicotinamide amide contact Asp964 (Asn240) and Thr929 (Thr209), respectively, the phosphates interact with Asp964 and Val965 (Val241), while the nicotinamide ribose hydrogen bonds Gly1075 (Ala321) and Asn1115 (Asn347). On the other hand, the 2’-OH of the adenosine ribose, along with the 3’-OH, interacts with Asp960 (Phe238) and is additionally surrounded by Asp983 (Gly261) and His984, occupying a relatively more spacious pocket than in *Cs*GDH ^8^. Considering that His984 and Arg985 (Asp263) might stabilize the 2’-phosphate of NADP^+^, one possible explanation for mL-GDH_180_ specificity for NAD^+^ ^10,11^ is that this enzyme, as well as related L-GDHs ^2,12–17^, evolved to favor NAD^+^ via the interactions of the 2’-OH group with Asp960 and Asp983. These residues might also exert a repulsive effect on the 2’-phosphate of NADP^+^. This interpretation aligns with previous suggestions that NAD^+^ specificity is determined by interactions between the 2’-OH of the adenosine ribose and both the Gly-rich loop (GxGxxA/S, as defined in L-GDHs_180_ ^2^) and an acidic residue at the so-called position 7 (Asp983 in mL-GDH_180_) of the fingerprint region of the nucleotide binding fold ^18,19^.

A seminal study on *Cs*GDH established that amino acid specificity is conferred by Lys89 and Ser380, which interact with the γ-carboxylate of the substrate ^20^. In mL-GDH_180_, Arg780 (Lys89) is suitably positioned to form salt bridges with the γ-carboxylate of L-glutamate (Figure 2). Though the hydroxyl of Ser1147 (Ser380) lies 4.3 Å from the γ-carboxylate of L-glutamate, slightly beyond hydrogen bond range, its position corresponds to that of Ser360 in *Cs*GDH. Also, residues Ala886 and Val1144 (Ala163 and Val377), Gly814 and Ala815 (Gly122 and Gly123), Gly781 and Gly782 (Gly90 and Gly91) and Gly1143 (Gly376), previously highlighted for their roles in establishing hydrophobic interactions with the γ- and δ-carbons of L-glutamate, shaping the L-glutamate binding pocket, orienting functional groups in the catalytic center and being located near the binding site of the nicotinamide ring of the dinucleotide, respectively ^20^, are similarly arranged in the mL-GDH_180_ cofactor/ligand-bound monomers. This structural similitudes underscore the conservation of key amino acid specificity determinants in mL-GDH_180_ (and related L-GDHs; see Figure S5 and ^2^) and S-GDHs.

Our steady-state data support an ordered sequential mechanism for the oxidative deamination of L-glutamate catalyzed by mL-GDH_180_, in which NAD⁺ binds first to the active site, followed by L-glutamate; chemistry then ensues, 2-oxoglutarate is released first, and NADH leaves last (Figure S1D and S1E). In line with this, our cryo-EM 3D classifications recovered empty, cofactor and cofactor/ligand monomer species but not a distinct ligand-only class of subunits. Moreover, given the conservation of residues that mediate catalysis, L-GDHs are expected to function similarly to S-GDHs ^3,20^. Our atomic model of the mL-GDH_180_ cofactor/ligand-monomer aimed to account for the density observed in the GluBR (Figure 2). In this model, the α-carboxylate and amino groups of L-glutamate adopt a slightly different orientation than in *Cs*GDH ^20^. Although the α-carboxylate of L-glutamate faces Lys806 (Lys113), their separation (4.1 Å) is longer than ideal for direct interaction. Similarly, the α-carboxylate of the substrate lies 6.4 Å from the side chain of Gln803 (Gln110), and the amino group is 4.9 Å from the carbonyl of Ala887 (Gly164) and 6.7 Å from the side chain of the catalytic Asp888 (Asp165). It is worth noting that, at our working resolution, we cannot detect water molecules that might contribute to substrate stabilization within the active site. Besides, the packing of residues 825-838 against segment 889-894 may help stabilize the position of Asp888, even though its side chain is only partially defined in cryo-EM maps. Certainly, catalysis is expected to involve a localized conformational adjustment that enables the side chain of Asp888 to approach the substrate’s amino group for proton abstraction in the first step of the reaction ^20^. Nevertheless, the cofactor/ligand-monomer maps (Figure 2 and Figure 4) reveal density corresponding to a group modeled as the α-carbon of L-glutamate, located 3.2 Å from the C4 atom of the cofactor nicotinamide, and density 3-3.5 Å from the Ala887 carbonyl, consistent with the position of the leaving NH_3_ group during catalysis ^20^.

Taken together with previous findings ^3^, our data confirm that mL-GDH_180_ exhibits pronounced conformational flexibility. In the absence of ligands, the enzyme populates two tetrameric forms, Open and Closed, detected at 4.19 and 6.6 Å resolution, respectively ^3^. In the present study, under conditions of saturating NAD^+^ (4 mM) and sub-saturating L-glutamate (42 mM) (see also Figure S1), three tetrameric states, Open, Closed1 and Closed2, were observed, with evidence of DNBR occupancy across the full ensemble (Figure 1). Thus, compared to the ligand-free state, these conditions appear to shift the conformational equilibrium toward the Closed forms. The segment 825-838 adopts distinct geometries depending on whether the cofactor is anchored to the DNBR and whether L-glutamate (or derivatives thereof) is present in the GluBR (Figure 2). In monomers where both regions are simultaneously occupied (cofactor/ligand-monomers), the conformation of this segment is compatible with an interaction with residues 704-706 of the N-terminally linked neighboring subunit in Closed tetramers (Figure 4).

Unlike in S-GDH ^20,21^, we did not observe changes in the relative positions of SDI and SDII within the catalytic domain of mL-GDH_180_. Instead, conformational transitions among oligomeric states were accompanied by adjustments at the ACT*3 and C-terminal domain interfaces, changes in the intersubunit contact between residues 650 (in the HM3) and 790 (in the catalytic domain), and the linkage between segments 825-838 and 704-706 (both in the catalytic domain), the latter forming only in Closed tetramers when both substrate and cofactor occupy the active site (Figure 5). These long-range transitions are presumably slower than catalysis and are thus observable in our experiment. While our data do not permit conclusions about positional and/or compositional heterogeneity within the GluBR or DNBR during catalysis, the most reasonable scenario is that the reaction occurs when both regions are loaded (Figure 4), with the cofactor approaching the substrate binding pocket (Figure 2). Overall, our results indicate that the 825-838/704-706 bridge acts as a closure-dependent modulator that biases the co-occupied Open-Closed equilibrium and tunes the dwell time of the Closed ternary complex; the net effect on turnover is expected to reflect the kinetic balance between chemistry and product release under the conditions considered.

**Figure 5.**
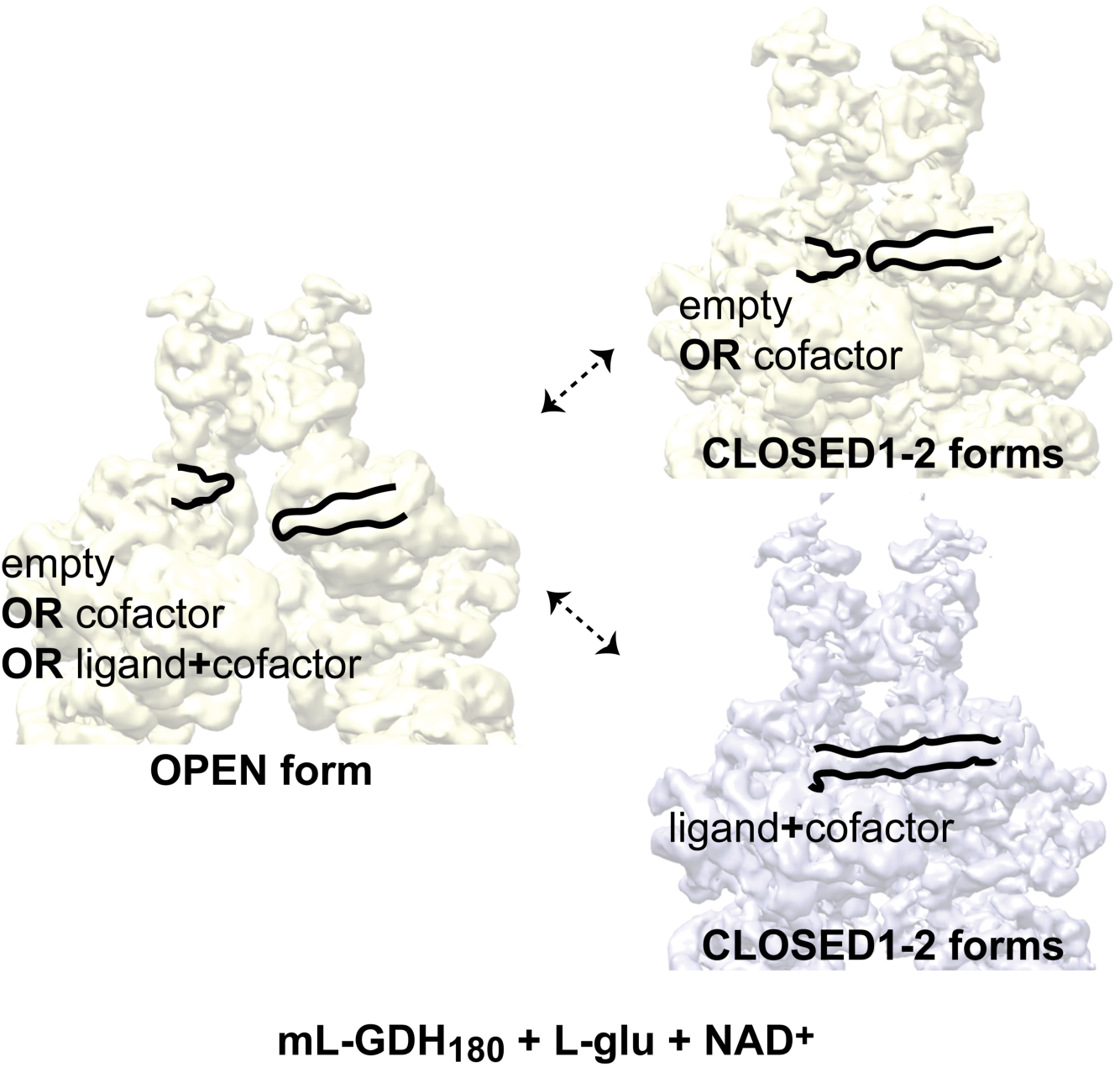
Ligand binding triggers tertiary structure remodeling that links catalysis with intersubunit communication in mL-GDH_180_. Cryo-EM maps of mL-GDH₁₈₀ incubated with L-glutamate and NAD⁺ illustrate the conformational ensemble comprising one open and two closed tetrameric states. In the open form, active sites can be empty or occupied by the cofactor or by both cofactor and ligand. In the closed conformations, when both the cofactor and the substrate simultaneously occupy the active site, the catalytic domains of neighboring subunits connected through their N-terminal regions approach each other, forming an intersubunit contact approximately 2 nm away from the catalytic center bearing the ligand and cofactor. This structural transition provides a physical link between active-site occupancy and long-range communication within the tetramer, coupling catalysis to quaternary-structure remodeling.

Notably, in the absence of added effectors, we found no evidence of cooperativity for mL-GDH_180_ catalyzed L-glutamate oxidation across pH 6.0–7.1, an interval that spans the reaction’s apparent pH optimum (Figure S1B) and overlaps the physiological pH range ^22,23^. At pH 6.8, the initial-rate dependences on L-glutamate and on NAD⁺ were Michaelis–Menten both when the co-substrate was fixed at a saturating concentration and when it was held at several subsaturating levels (Figure S1D). Consistent with this trend, datasets acquired at pH 6.5 (the condition used for cryo-EM specimen preparation) were likewise well described by Michaelis–Menten kinetics. Thus, within this near-optimal pH window, steady-state behavior is consistent with independent active sites. The structural origin of departures from this pattern at basic pH (this work and ^10^) and under activity modulators ^3,10^ remains to be determined, particularly given the regulatory potential of the N- and C-terminal regions of the enzyme.

We found that ligand binding to the active (orthosteric) site of mL-GDH_180_ is linked to tertiary structure changes that lead to the consolidation of an oligomeric interface two nanometers away from the catalytic center ^24^. It is worth considering whether and how natural or designed molecules binding to the 825-838 and/or 704-706 segments could influence catalysis and whether occupancy of the active site modulates such regulation ^25^. Our findings open new avenues for high-resolution studies on catalysis and regulation of L-GDHs of medical and industrial relevance.

## MATERIALS AND METHODS

### Protein production

Plasmid pLIC-His-mL-GDH_180_ ^3^ was used to produce N-terminally His6-tagged mL-GDH180 (MSMEG_4699, Uniprot A0R1C2) in *E. coli* cells. Transformed *E. coli* BL21(DE3) cells were grown at 37 °C in LB broth supplemented with 100 µg/ml ampicillin until reaching 0.8 units of optical density at 600 nm. Protein expression was then induced by adding isopropyl β-D-1-thiogalactopyranoside (IPTG) to a final concentration of 0.5 mM and incubation was continued for 18 hours at 14 °C. The culture was centrifuged and the pellet was resuspended in 25 mM Hepes, 500 mM NaCl, 10 % V/V glycerol, 10 mM imidazole, pH 8.0. The suspension was supplemented with 0.01 µg/ml DNAse, 4 µM MgSO_4_ and the protease inhibitor cOmplete™ EDTA-free Protease Inhibitor Cocktail (Roche). Cells were then sonicated and the mixture was clarified by centrifugation. The supernatant was loaded onto a HisTrap HP column (Cytiva) equilibrated with buffer 25 mM Hepes, 500 mM NaCl, 10% v/v glycerol, 20 mM imidazole, pH 8.0, and His6-tagged mL-GDH_180_ was purified by applying a linear imidazole gradient (20-500 mM). The protein was then further purified by size-exclusion chromatography, using a Superose 6 10/300 GL column (Cytiva) equilibrated in 20 mM Mes, 300 mM NaCl, pH 6.5. mL-GDH_180_ containing fractions, as confirmed by SDS-PAGE, were pooled and used immediately or stored at -80 °C for use in subsequent studies. The protein was quantified by electronic absorption using the molar absorption coefficient of 171,090 M^-1^ cm^-1^, predicted from the amino acid sequence by the ProtParam tool (http://web.expasy.org/protparam/).

### Steady state kinetics

Reactions were performed at 25 °C, using 10 nM mL-GDH_180_. Oxidative deamination of L-glutamate catalyzed by mL-GDH_180_ was monitored by the increase in absorbance at 340 nm due to NADH production (ε_340_ = 6220 M^-1^ cm^-1^) on a Shimadzu UV-120-02 spectrophotometer. Initial rate measurements were carried out in the following buffers: 50 mM Mes, 50 mM NaCl, pH 6.0; 50 mM Mes, 50 mM NaCl, pH 6.5; 50 mM Hepes, 50 mM NaCl, pH 6.8; 50 mM Hepes, 50 mM NaCl, pH 7.5; 50 mM Hepes, 50 mM NaCl, pH 8.2; 30 mM sodium phosphate buffer, pH 7.1; 30 mM sodium phosphate buffer, pH 8.0. Within pH 6.0–7.1, the kinetic parameters *k*_cat_ and *K*_M_ were obtained by fitting the dependence of the initial rate on substrate concentration to the Michaelis-Menten equation (Figure S1). At pH 7.5, the dependence of the initial rate on substrate concentration was sigmoidal; accordingly, data were fitted to the Hill equation. Parameter estimation for Michaelis-Menten and Hill models was performed by nonlinear regression. At pH 8.0 and 8.2, where activity was low and the rate-substrate relationship remained essentially linear across the full substrate range, catalytic efficiency (*k*_cat_/*K*_M_) was obtained by linear regression of rate versus substrate concentration. Each experiment was performed at least twice, in some cases using independent protein preparations. When error bars are shown on kinetic parameters, they represent the standard errors from curve fitting. Mean *k*_cat_ and *K*_M_ values reported in the text include errors obtained by propagating the standard errors from individual fits across replicates; catalytic efficiency was calculated as *k*_cat_/*K*_M_ with error propagated from those parameter errors.

### Cryo-electron microscopy

4 µl of a mixture containing 42 mM L-glutamate, 4 mM NAD^+^ and 0.125 mg/ml mL-GDH_180_ was prepared in 20 mM Mes, 300 mM NaCl, pH 6.5, at 10 °C, using a buffer closely aligned with conditions previously optimized for single-particle analysis ^3^. Samples were applied to Quantifoil R2/2 holey carbon grids and vitrified using a Leica EM GP2 within the first minute after mixing, during which enzymatic activity was confirmed by an increase in absorbance at 340 nm (Figure S1C), indicative of NADH production. In addition, a similar protein solution was applied onto the same type of grids bearing an additional thin layer of carbon. Vitrification on these two types of grids was required to obtain top and side views of mL-GDH_180_ tetramers (Figure S2). Data collection was carried out in a ThermoFisher Titan Krios G4 transmission electron microscope paired with a Gatan BioContinuum Imaging Filter and a K3 direct electron detector, at the BREM (Basque Resource for Electron Microscopy). 23,747 and 19,038 movie frames were acquired for the sample on R2/2 grids and grids supplemented with an additional thin layer of carbon, respectively (Figure S2). Both collections were performed at a nominal magnification of ×130,000, resulting in a sampling of 0.6462 Å/pixel. Each movie contained 50 frames with an accumulated dose of 49 e^-^/Å^2^. Movie frames were aligned using MotionCor2 ^26^ and the final average included all dose-weighted frames.

The contrast transfer function (CTF) of the micrographs was estimated using CTFFIND4 ^27^. Particles were automatically selected from the micrographs using autopicking from RELION-4 ^28^. To speed up calculations, particles were extracted with a 2x binning, resulting in a pixel size of 1.29 Å/pixel. In datasets with potential for high resolution, particles were re-extracted at original sampling. An initial set of 667,866 particles was subjected to 2D and 3D classification procedures (Figure S3). The 3D-classification with imposed D2 symmetry resulted in eight different classes, but only three achieved good definition and high resolution. Three different conformations were obtained, called the Open (37,641 particles), Closed1 (45,998 particles) and Closed2 (32,441 particles) conformations, with estimated resolutions of 3.88 Å, 3.54 Å and 3.57 Å, respectively. Each class was symmetry-expanded, the star files were joined (116,080×4, *i.e.* 464,320 particles) and an average 3D map of a subunit was refined using a mask and polished to a final resolution of 2.63 Å. A 3D classification within a sphere mask focused on the active site separated three different states: empty-monomer, with 131,216 particles and 3.07 Å of resolution, cofactor-monomer, with 168,668 particles and 2.95 Å of resolution, and cofactor/ligand-monomer, with 112,701 particles and 3.12 Å of resolution. These three groups were crossed (common particles between selection files) with the three sets of particles obtained in the initial classification from the conformational changes of the tetramer, resulting in nine groups of particles that were refined independently (Figure S3). The three maps with signal in the cofactor and substrate binding pockets were subjected to polishing and ctfrefine to improve the resolution, each one coming from a conformation of the tetramer: Open-cofactor/ligand-monomer (3.23 Å), Closed1-cofactor/ligand-monomer (3.11 Å) and Closed2-cofactor/ligand-monomer (3.18 Å). The resolution of the cryo-EM density maps was estimated using a threshold of 0.5 in the Fourier Shell Correlation (FSC) between independently processed halfmaps, *i.e.*, gold standard. The cryo-EM density maps were postprocessed by local sharpening using EMReady ^29^, which improved the structural details and facilitated the atomic model building.

Model fitting into cryo-EM maps was performed using the programs UCSF Chimera ^30^, Coot ^31^ and phenix.real_space_refine ^32^. From the 2.63 Å resolution cryo-EM map of a mL-GDH_180_ monomer and the atomic coordinates from the Protein Data Bank entry 7A1D, a model of the polypeptide chain was refined which was then fitted into the other cryo-EM maps. After removing parts of the model not defined in the maps, performing manual model building when necessary and placing ligands manually, using Coot ^31^, where appropriate, atomic coordinates and B-factors were refined using phenix.real_space_refine ^32^ with NCS and secondary structure restraints. RMSDs between models were calculated using Coot ^31^. Figures were generated and rendered with UCSF Chimera ^30^ or PyMOL version 1.8.x (Schrödinger, LLC).

Cryo-EM maps obtained for mL-GDH_180_ were deposited in the Electron Microscopy Data Bank under the accession codes EMD-52419 (Open), EMD-52420 (Closed1), EMD-52421 (Closed2), EMD-52422 (empty-monomer), EMD-52423 (cofactor-monomer), EMD-52424 (cofactor/ligand-monomer), EMD-52425 (Open-cofactor/ligand-monomer), EMD-52426 (Closed1-cofactor/ligand-monomer), EMD-52427 (Closed2-cofactor/ligand-monomer), EMD-52428 (total-monomer) and EMD-52429 (Closed2-cofactor/ligand-tetramer). Atomic coordinates were deposited in the Protein Data Bank under the accession codes 9HUX (Open), 9HUY (Closed1), 9HUZ (Closed2), 9HV0 (empty-monomer), 9HV4 (cofactor-monomer), 9HV5 (cofactor/ligand-monomer) and 9HV6 (total-monomer).

Description of supplementary material including filenames

Supplementary material.doc: Supplementary figures and tables.

Movie S1.mp4: Changes in the quaternary structure of mL-GDH_180_ detected by cryo-EM in the presence of L-glutamate and NAD^+^.

Movie S2.mp4: Changes in the active site of mL-GDH_180_ detected by cryo-EM in the presence of L-glutamate and NAD^+^.

Movie S3.mp4: In the presence of L-glutamate and NAD^+^, a contact is established between the catalytic domains of neighboring mL-GDH_180_ subunits connected through their N-terminal regions.

## ACKNOWLEDGMENTS

This work was supported by grants PICT 2019-00989 and PICT 2021-00253 from the Agencia Nacional de Promoción de la Investigación, el Desarrollo Tecnológico y la Innovación (Agencia I+D+i, Argentina) to MNL as well as grants PID2021-124074NB-I00 and PID2021-125946OB-I00 from Agencia Estatal de Investigación of Spain to MV and GJO, respectively. We also thank the Severo Ochoa Center of Excellence Accreditation CEX2021-001136-S. MNL is a career investigator from Consejo Nacional de Investigaciones Científicas y Técnicas (CONICET, Argentina). NC holds a CONICET PhD fellowship. NC and MNL are grateful to Liliana Rojas for excellent technical assistance. Some of this work was performed at the Basque Resource for Electron Microscopy located at Instituto Biofisika (UPV/EHU, CSIC), supported by the Department of Science, Universities and Innovation and the Innovation Fund of the Basque Government, with additional support from MCIN (Recovery, Transformation and Resilience Plan) and the Basque Government “Biotechnology Complementary Plan Applied to Health” with funding from European Union NextGenerationEU (PRTR-C17.I1; PRTR-C17.I01.P01.S13) (AAAA_ACG_AY_2539/22_05).

## AUTHOR CONTRIBUTIONS

ML produced protein, prepared EM samples, performed EM experiments, processed and analyzed EM data and performed structural analyses; NC produced protein, performed kinetic analyses, analyzed EM data and performed structural analyses; JPLA collected and analyzed EM data; DC assisted during processing of EM data; RMR performed structural analyses; GJO performed structural analyses; MV designed experiments, supervised EM data collection and processing and analyzed results; MNL designed experiments, supervised biochemical experiments, refined atomic models, analyzed data and wrote the paper. All authors read and corrected the paper.

