## Supplemental Data for "Tertiary and quaternary structure remodeling by occupancy of the substrate binding pocket in a large glutamate dehydrogenase"

Running title: Ligand-driven remodeling of mL-GDH$_{180}$

**SUPPLEMENTARY MATERIAL**

**SUPPLEMENTARY FIGURES**

**Figure S1**

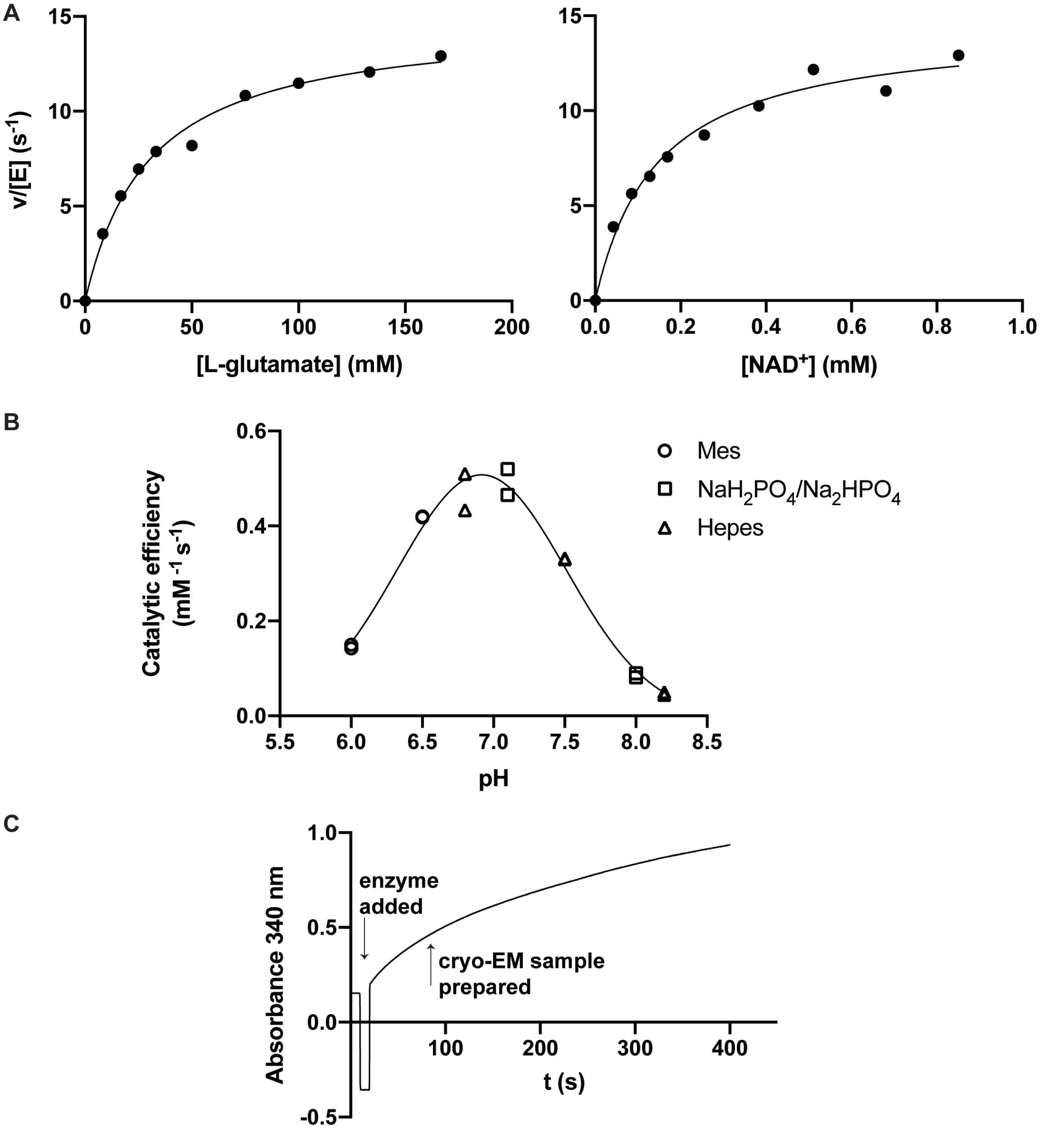

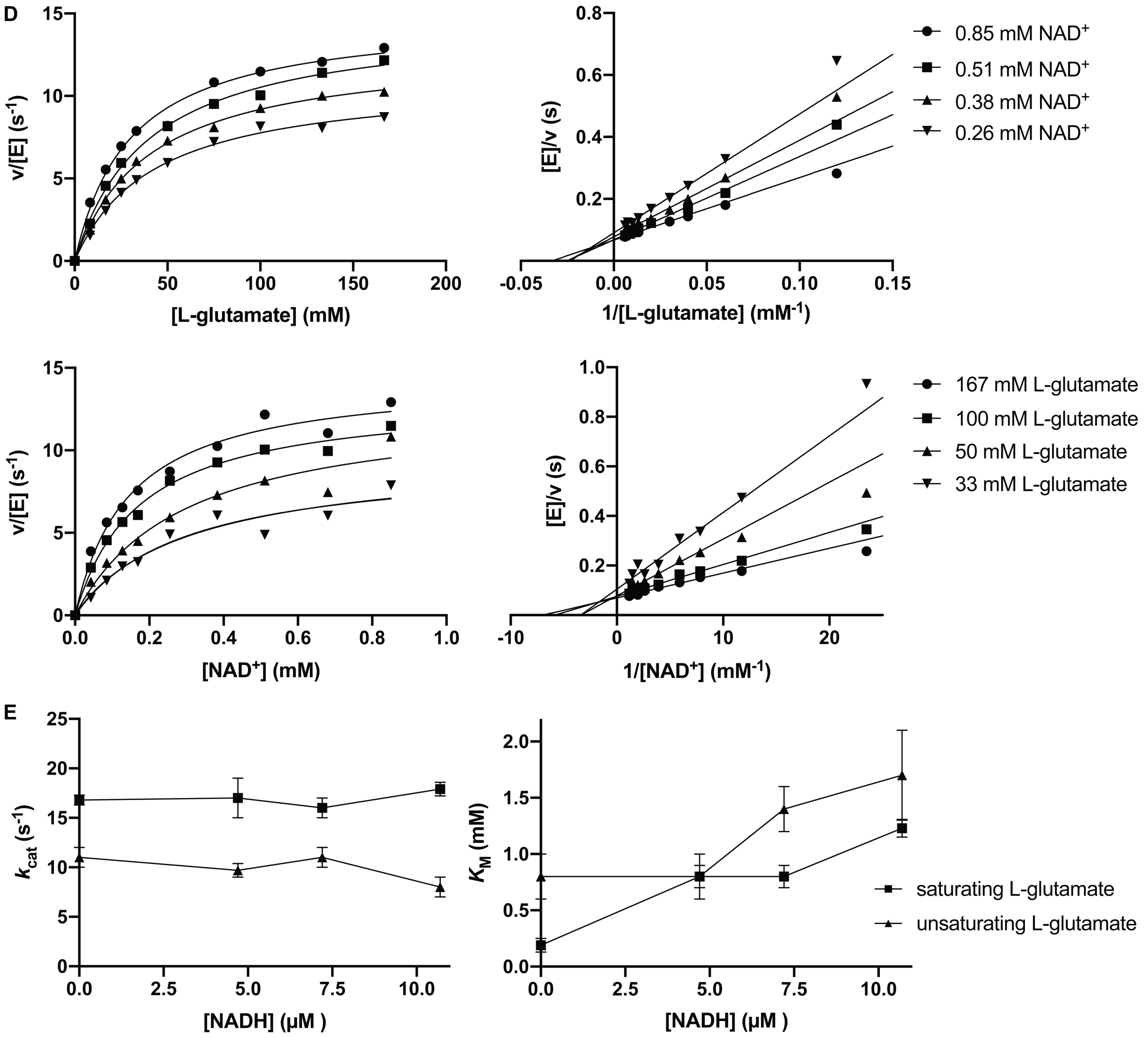


**Oxidative deamination of L-glutamate by mL-GDH$_{180}$: biochemical characterization.** (A) Representative initial rate dependences on L-glutamate (left) and on NAD$^+$ (right) in 50 mM Hepes, 50 mM NaCl, pH 6.8. The co-substrate was held at saturating concentration (1 mM NAD$^+$ for the L-glutamate series; 170 mM L-glutamate for the NAD$^+$ series); solid curves are Michaelis-Menten fits obtained by nonlinear regression. (B) pH dependence of the catalytic efficiency ($k_{cat}/K_M$) derived from initial rate data with NAD$^+$ at 1 mM; assays were carried out in Mes, sodium phosphate or Hepes buffers as detailed in Materials and methods. Values of $k_{cat}/K_M$ were obtained from the fits described therein: Michaelis-Menten (nonlinear regression) across pH 6.0-7.1, Hill equation at pH 7.5 (sigmoidal behavior; Hill coefficient ≈ 2) and linear regression of rate versus substrate concentration at pH 8.0-8.2, where the response was essentially linear. Each symbol corresponds to $k_{cat}/K_M$ from a single independent dataset. The solid line is a Gaussian fit used to estimate the apparent pH optimum, which peaks just below neutrality. (C) Representative progress curve at 340 nm showing NADH formation under the solution composition used for cryo-EM sample preparation (42 mM L-glutamate, 4 mM $NAD^+$ and 0.125 mg/ml mL-$GDH_{180}$ in 20 mM Mes, 300 mM NaCl, pH 6.5). The downward arrow indicates the moment of enzyme addition; the upward arrow marks the time point at which the grid was vitrified. (D) Top row: left, initial rate versus L-glutamate at fixed $NAD^+$ concentrations; right, the corresponding double-reciprocal (Lineweaver-Burk) plots. Bottom row: left, initial rate versus $NAD^+$ at fixed L-glutamate concentrations; right, the corresponding double-reciprocal plots. Solid curves in the left panels are Michaelis-Menten fits obtained by nonlinear regression. Right-panel Lineweaver-Burk plots were generated by reciprocal transformation of the experimental data points and the best-fit Michaelis-Menten functions. Assays were carried out in 50 mM Hepes, 50 mM NaCl, pH 6.8. The non-parallel lines that intersect in the second quadrant are indicative of a sequential mechanism (rather than ping-pong). (E) Effect of added NADH on kinetic parameters derived from initial rate fits to $NAD^+$-dependence curves ($NAD^+$ varied; L-glutamate fixed at 170 mM or 30 mM) in 50 mM Hepes, 50 mM NaCl, pH 6.8. NADH increased the apparent $K_M$ for $NAD^+$ with little or no effect on $k_{cat}$, consistent with competitive product inhibition versus $NAD^+$ and an ordered sequential mechanism in which $NAD^+$ binds first and NADH is released last. Solid lines are guides to the eye.

**Figure S2**

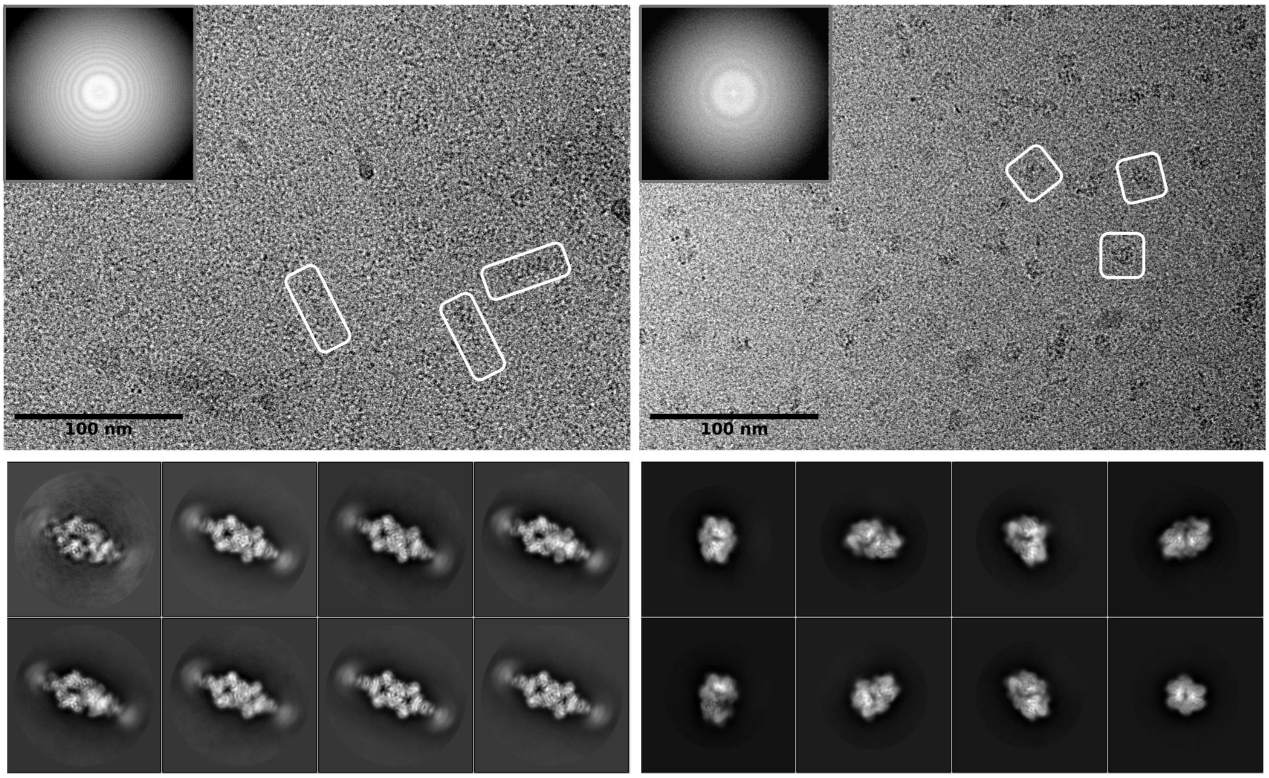


**Cryo-EM of mL-GDH$_{180}$ on two grid types.** The top-left panel shows a representative micrograph of the sample on a grid with a thin layer of carbon as additional support, together with a calculated power spectrum. Some single particles attributed to side views of tetrameric mL-GDH$_{180}$ are indicated with boxes. The bottom-left panel includes some 2D averages for particles extracted from micrographs as in top-left panel. The top-right panel depicts a micrograph of mL-GDH$_{180}$ tetramers on holey carbon grids where some top views for particles are highlighted with boxes. The calculated power spectrum for this micrograph is also included. The bottom-right panel shows 2D averages obtained for particles coming from data as in top-right panel.

#### Figure S3

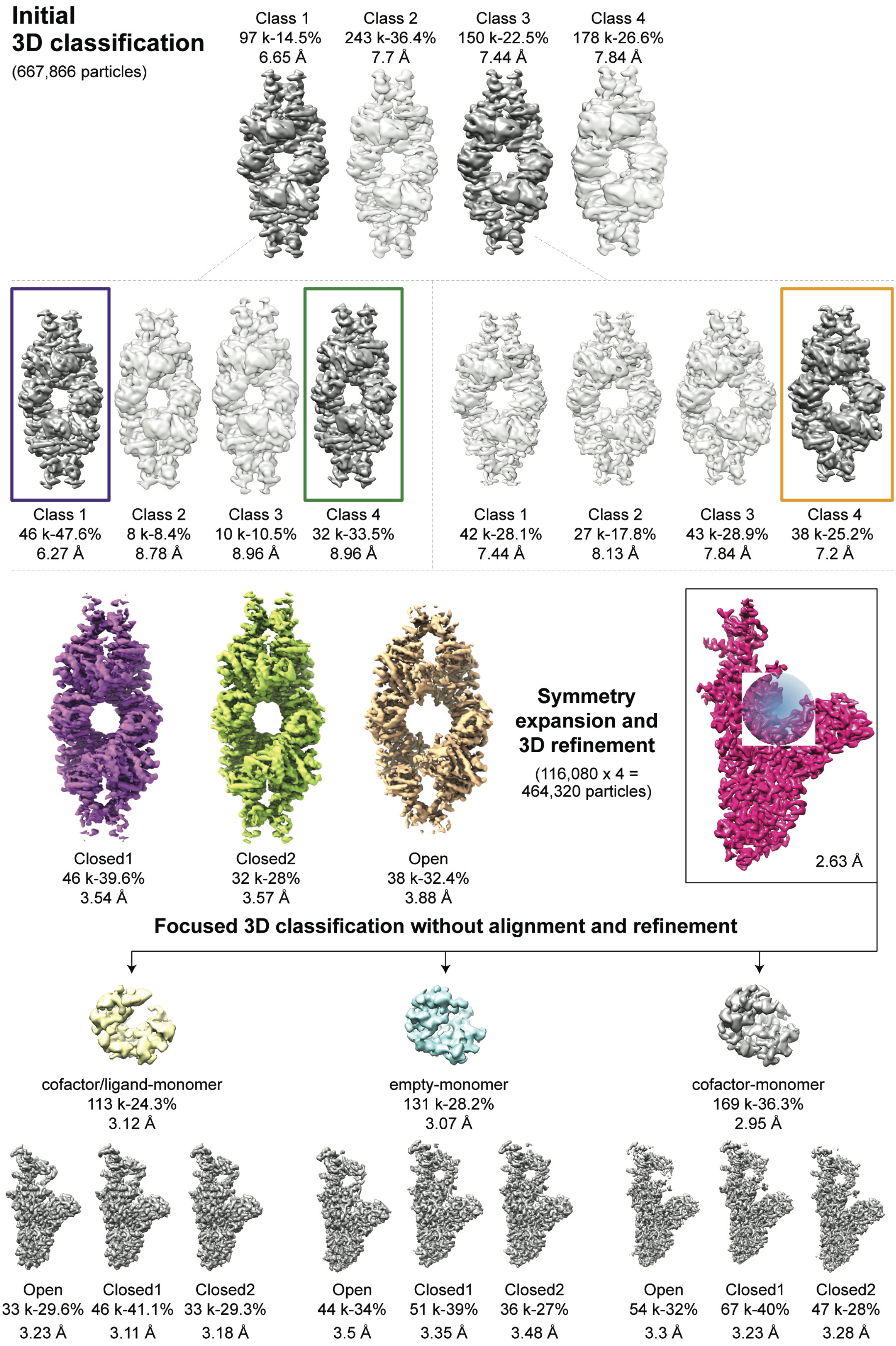

Initial
3D classification
(667,866 particles)
Class 1
97 k-14.5%
6.65 Å
Class 2
243 k-36.4%
7.7 Å
Class 3
150 k-22.5%
7.44 Å
Class 4
178 k-26.6%
7.84 Å
Class 1
46 k-47.6%
6.27 Å
Class 2
8 k-8.4%
8.78 Å
Class 3
10 k-10.5%
8.96 Å
Class 4
32 k-33.5%
8.96 Å
Class 1
42 k-28.1%
7.44 Å
Class 2
27 k-17.8%
8.13 Å
Class 3
43 k-28.9%
7.84 Å
Class 4
38 k-25.2%
7.2 Å
Closed1
46 k-39.6%
3.54 Å
Closed2
32 k-28%
3.57 Å
Open
38 k-32.4%
3.88 Å
Symmetry expansion and 3D refinement
(116,080 x 4 = 464,320 particles)
2.63 Å
Focused 3D classification without alignment and refinement
cofactor/ligand-monomer
113 k-24.3%
3.12 Å
empty-monomer
131 k-28.2%
3.07 Å
cofactor-monomer
169 k-36.3%
2.95 Å
Open
33 k-29.6%
3.23 Å
Closed1
46 k-41.1%
3.11 Å
Closed2
33 k-29.3%
3.18 Å
Open
44 k-34%
3.5 Å
Closed1
51 k-39%
3.35 Å
Closed2
36 k-27%
3.48 Å
Open
54 k-32%
3.3 Å
Closed1
67 k-40%
3.23 Å
Closed2
47 k-28%
3.28 Å

**Cryo-EM data processing.** Scheme of the particle classification procedure performed to obtain the final maps corresponding to the different conformations of mL-$GDH_{180}$ detected upon incubation with L-glutamate and $NAD^+$, including refinement, symmetry expansion for monomer processing and classification with a sphere mask centered on the active site. For each map, the number of particles and the resolution achieved are indicated.

**FIGURE S4**

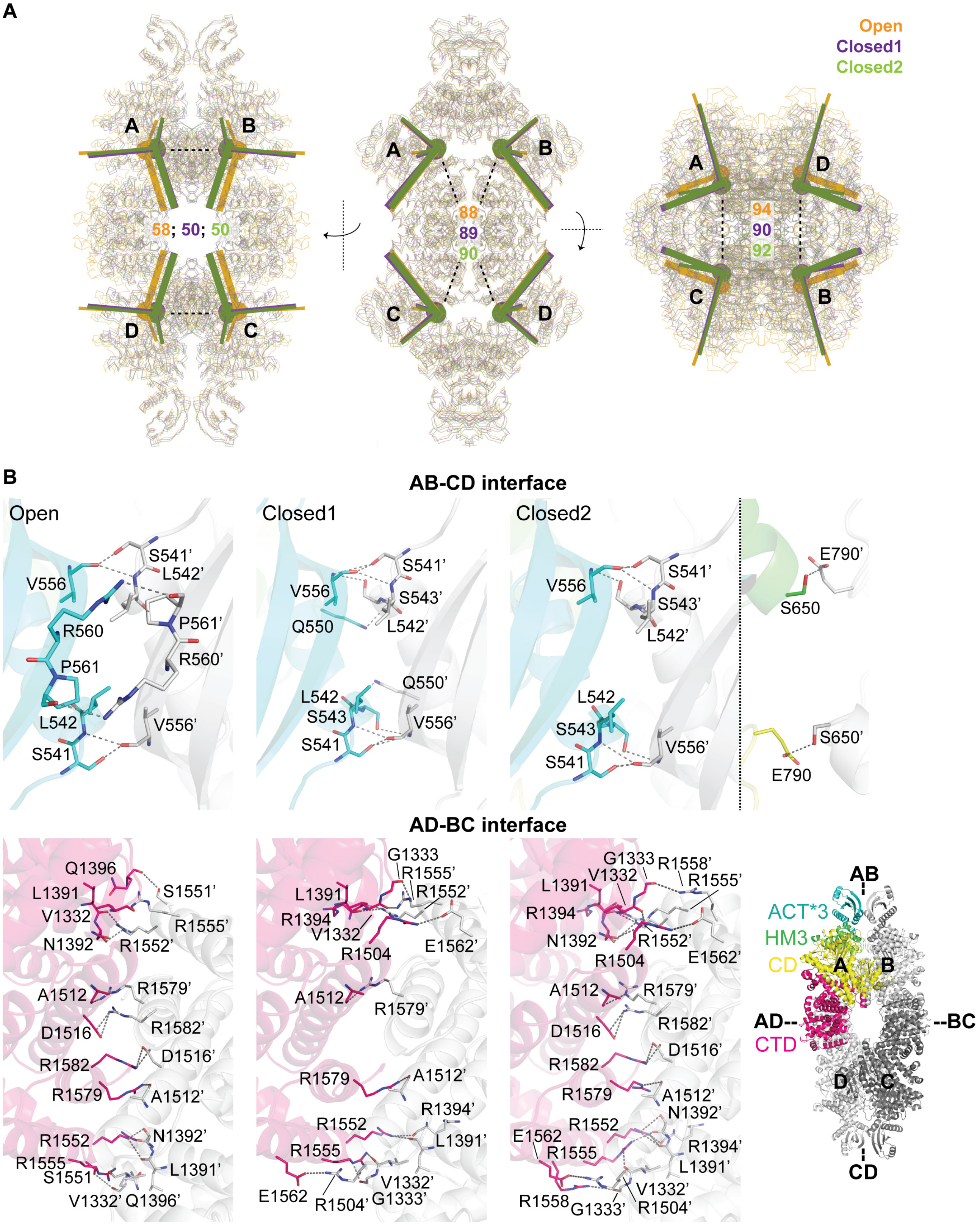


**Rearrangements of mL-GDH$_{180}$ quaternary structure.** (A) The centers of mass (COMs; circles) and inertia tensors (ITs; lines) of each subunit in the three conformers are shown from the front, side and top of the whole tetramers. COMs and ITs were calculated from the positions of all α-carbons. While the orientation of each subunit varies only slightly between conformations, there is a significant shift in the position of the COMs when going from the Open to Closed conformations. The Open, Closed1 and Closed2 tetramers are shown in orange, violet and green, respectively. The distances between the COMs of monomers **A**, **B**, **C** and **D** are indicated with dashed lines and are expressed in Å. (B) Intersubunit interactions in the different mL-GDH$_{180}$ conformers. Atomic models are shown in ribbon representation, with interacting residues depicted as sticks. Domains are colored for monomer **A** following the reference provided in the model in the lower right corner, where the position of the AD-BC and AB-CD interfaces in the complex is also shown. At the AB-CD interface, the side chain (sc) of Ser541 and the main chain (mc) of Leu542 and Val556 are involved in interfacial hydrogen bonds in all three mL-GDH$_{180}$ tetrameric forms. Besides, Arg560 (sc) hydrogen bonds Pro561 (mc) exclusively in the Open conformation, Ser543 (mc) interacts with Gln550 (sc) only in the Closed1 form, while Ser543 (sc) also hydrogen bonds Val556 (mc) in both the Closed1 and Closed2 mL-GDH$_{180}$ conformers. Residues Val556, Arg560 and Pro561 are located in strand β3 (554-565) of the ACT*3 domain, while Ser541, Leu542, Ser543 and Gln550 are situated at the edges of its helix α1 (541-550). On the other hand, as a result of contacts at the AD-BC interface, in the Open form of the enzyme, helix 1551-1566 of one monomer interacts simultaneously with helix 1324-1332, helix 1369-1393 and loop 1394-1399 of the adjacent one, and helix 1571-1583 of one subunit contacts helix 1504-1530 of the other. In the Closed1 conformer, helix 1551-1566 additionally interacts with loop 1333-1336 and helix 1504-1530. Compared to the Closed1 form, the Closed2 conformer establishes an alternative series of contacts, although they involve the same set of secondary structure elements. The ACT*3 domain (residues 513-617), helical motif 3 (HM3; residues 618-701), catalytic domain (CD; residues 702-1220) and C-terminal domain (CTD; residues 1221-1594) of mL-GDH$_{180}$ were previously defined [1].

### Figure S5

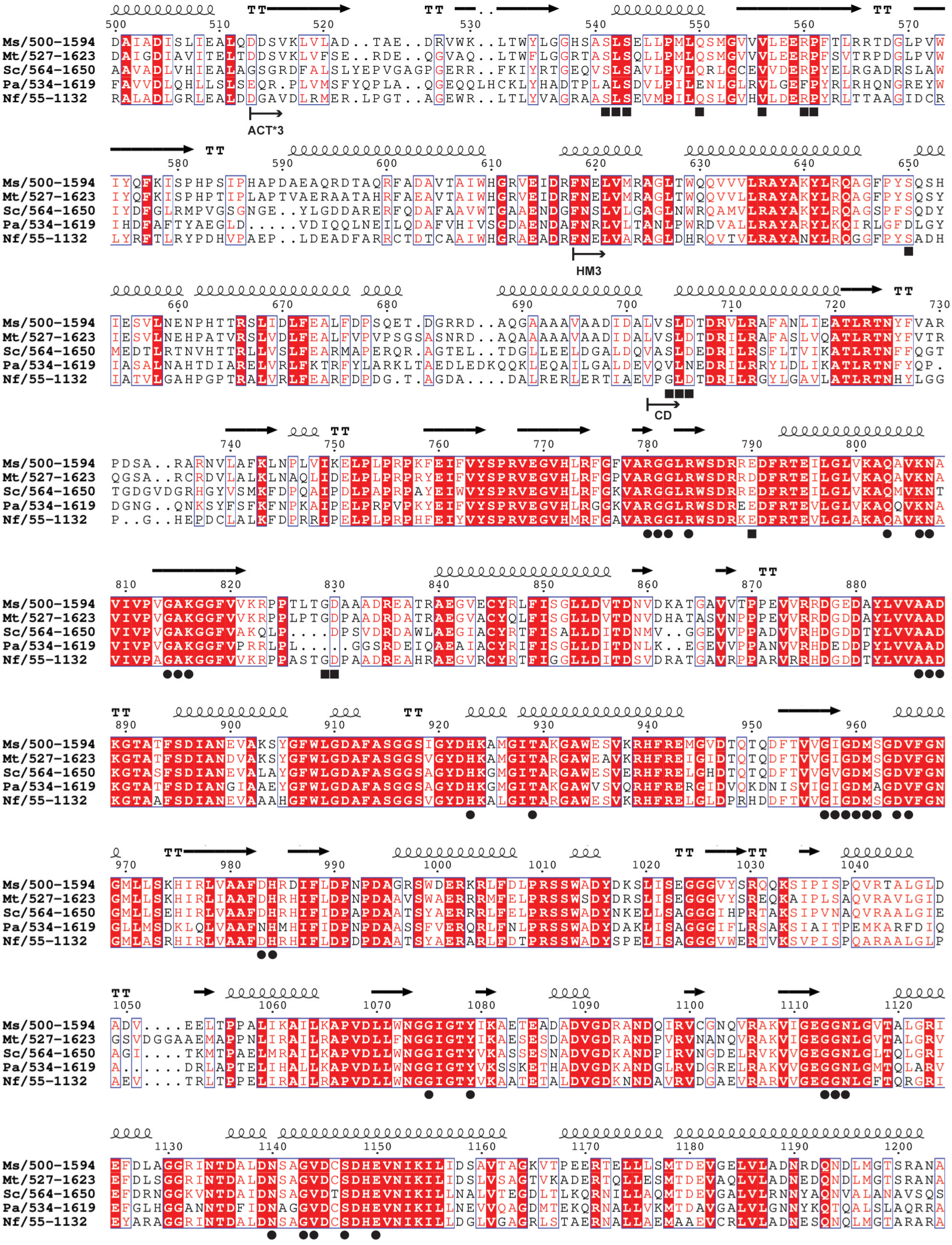

Ms/500-1594 DAIADISLIEALQDDSVKLVLAD..TAE..DRVWK..LTWYLGGHSASLSELLPMLQSMGVVVLEERPFTLRRTDGLPVW
Mt/527-1623 DAIGDIAVITELTDDSVKLVFSE..RDE..QGVAQ..LTWFLGGRTASLSQLLPMLQSMGVVVLEERPFSVTRPDGLPVW
Sc/564-1650 AAVADLVHIEALAGSGRDFALSLYEPVGAGPGERR..FKIYRTGEQVSLSAVLPVLQRLGCEVVDERPYELRGADRSLAW
Pa/534-1619 FAVVDLQHLLSLSEQR.PLVMSFYQPLA..QGEQQLHCKLYHADTPLALSDVLPILENLGLRVLGEFPYRLRHQNGREYW
Nf/55-1132 RALADLGRLEALDDGAVDLRMER.LPGT..AGEWR..LTLYVAGRAASLSEVMPILQSLGVHVLDERPYRLTTAAGIDCR
ACT*3
Ms/500-1594 IYQFKISPHPSIPHAPDAEAQRDTAQRFADAVTAIWHGRVEIDRFNELVMRAGLTWQQVVVLRAYAKYLRQAGFPYSQSH
Mt/527-1623 IYQFKISPHPTIPLAPTVAERAATAHRFAEAVTAIWHGRVEIDRFNELVMRAGLTWQQVVLLRAYAKYLRQAGFPYSQSY
Sc/564-1650 IYDFGLRMPVGSGNGE..YLGDDARERFQDAFAAVWTGAAENDGFNSLVLGAGLNWRQAMVLRAYAKYLRQAGSPFSQDY
Pa/534-1619 IHDFAFTYAEGLD.....VDIQQLNEILQDAFVHIVSGDAENDAFNRLVLTANLPWRDVALLRAYARYLKQIRLGFDLGY
Nf/55-1132 LYRFTLRYPDHVPAEP..LDEADFARRCTDTCAAIWHGRAEADRFNELVARAGLDHRQVTVLRAYANYLRQGGFPYSADH
HM3
Ms/500-1594 IESVLNENPHTTRSLIDLFEALFDPSQET.DGRRD..AQGAAAAVAADIDALVSLDTDRVLRAFANLIEATLRTNYFVAR
Mt/527-1623 IESVLNEHPATVRSLVDLFEALFVPVPSGSASNRD..AQAAAAAVAADIDALVSLDTDRILRAFASLVQATLRTNYFVTR
Sc/564-1650 MEDTLRTNVHTTRLLVSLFEARMAPERQR.AGTEL..TDGLLEELDGALDQVASLDEDRILRSFLTVIKATLRTNFFQGT
Pa/534-1619 IASALNAHTDIARELVRLFKTRFYLARKLTAEDLEDKQQKLEQAILGALDEVQVLNEDRILRRYLDLIKATLRTNFYQP.
Nf/55-1132 IATVLGAHPGPTRALVRLFEARFDPDG.T.AGDA....DALRERLERTIAEVPGLDTDRILRGYLGAVLATLRTNHYLGG
CD
Ms/500-1594 PDSA..RARNVLAFKLNPLVIKELPLPRPKFEIFVYSPRVEGVHLRFGFVARGGLRWSDRREDFRTEILGLVKAQAVKNA
Mt/527-1623 QGSA..RCRDVLALKLNAQLIDELPLPRPRYEIFVYSPRVEGVHLRFGPVARGGLRWSDRRDDFRTEILGLVKAQAVKNA
Sc/564-1650 TGDGVDGRHGYVSMKFDPQAIPDLPAPRPAYEIWVYSPRVEGVHLRFGKVARGGLRWSDRREDFRTEILGLVKAQMVKNT
Pa/534-1619 DGNG..QNKSYFSFKFNPKAIPELPRPVPKYEIFVYSPRVEGVHLRGGKVARGGLRWSDREEDFRTEVLGLVKAQQVKNA
Nf/55-1132 P..G..HEPDCLALKFDPRRIPELPLPRPHFEIYVYSPRVEGVHMRFGAVARGGLRWSDRKEDFRTEVLGLAKAQAVKNA
Ms/500-1594 VIVPVGAKGGFVVKRPPTLTGDAAADREATRAEGVECYRLFISGLLDVTDNVDKATGAVVTPPEVVRRDGEDAYLVVAAD
Mt/527-1623 VIVPVGAKGGFVVKRPPLPTGDPAADRDATRAEGVACYQLFISGLLDVTDNVDHATASVNPPEVVRRDGDDAYLVVAAD
Sc/564-1650 VIVPVGAKGGFVAKQLP....DPSVDRDAWLAEGIACYRTFISALLDITDNMV..GGEVVPPADVVRHDGDDTYLVVAAD
Pa/534-1619 VIVPVGAKGGFVPRRLPL....GGSRDEIQAEAIACYRIFISGLLDITDNLK..EGEVVPPANVVRHDEDDPYLVVAAD
Nf/55-1132 VIVPAGAKGGFVVKRPPASTGDPAADREAHRAEGVRCYRTFIGGLLDITDSVDRATGAVRPPARVRRHDGDDTYLVVAAD
Ms/500-1594 KGTATFSDIANEVAKSYGFWLGDAFASGGSIGYDHKAMGITAKGAWESVKRHFREMGVDTQTQDFTVVGIGDMSGDVFGN
Mt/527-1623 KGTATFSDIANDVAKSYGFWLGDAFASGGSVGYDHKAMGITARGAWEAVKRHFREIGIDTQTQDFTVVGIGDMSGDVFGN
Sc/564-1650 KGTASFSDIANEVALAYGFWLGDAFASGGSAGYDHKGMGITARGAWESVERHFRELGHDTQTQDFTVVGVGDMSGDVFGN
Pa/534-1619 KGTATFSDIANGIAAEYGFWLGDAFASGGSAGYDHKGMGITAKGAWVSVQRHFRERGIDVQKDNISVIGIGDMAGDVFGN
Nf/55-1132 KGTAAFSDIANEVAAAHGFWLGDAFASGGSVGYDHKALGITARGAWESVKRHFRELGLDPRHDDFTVVGIGDMSGDVFGN
Ms/500-1594 GMLLSKHIRLVAAFDHRDIFLDPNPDAGRSWDERKRLFDLPRSSWADYDKSLISEGGGVYSRQQKSIPISPQVRTALGLD
Mt/527-1623 GMLLSKHIRLIAAFDHRHIFLDPNPDAAVSWAERRRMFELPRSSWSDYDRSLISEGGGVYSREQKAIPLSAQVRAVLGID
Sc/564-1650 GMLLSEHIRLVAAFDHRHIFIDPAPDAATSYAERRRLFELPRSSWADYNKELLSAGGGIHPRTAKSIPVNAQVRAALGIE
Pa/534-1619 GLLMSDKLQLVAAFNHMHIFIDPNPDAASSFVERQRLFNLPRSSWADYDAKLISAGGGIFLRSAKSIAITPEMKARFDIQ
Nf/55-1132 GMLASRHIRLVAAFDHRHIFLDPDPDAATSYAERARLFDTPRSSWADYSPELISAGGGVWERTVKSVPISPQARAALGLP
Ms/500-1594 ADV....EELTPPALIKAILKAPVDLLWNGGIGTYIKAETEADADVGDRANDQIRVCGNQVRAKVIGEGGNLGVTALGRI
Mt/527-1623 GSVDGGAAEMAPPNLIRAILRAPVDLLFNGGIGTYVKAESESDADVGDKANDPIRVNANQVRAKVIGEGGNLGVTALGRV
Sc/564-1650 AGI....TKMTPAELMRAILKAPVDLLWNGGIGTYVKASSESNADVGDKANDPIRVNGDELRVKVVGEGGNLGLTQLGRI
Pa/534-1619 A......DRLAPTELIHALLKAPVDLLWNGGIGTYVKSSKETHADVGDKANDGLRVDGRELRAKVVGEGGNLGMTQLARV
Nf/55-1132 AEV....TRLTPPELIRAILRAPVDLLWNGGIGTYVKAATETALDVGDKNNDAVRVDGAEVRARVVGEGGNLGFTQRGRI
Ms/500-1594 EFDLAGGRINTDALDNSAGVDCSDHEVNIKILIDSAVTAGKVTPEERTELLLSMTDEVGELVLADNRDQNDLMGTSRANA
Mt/527-1623 EFDLSGGRINTDALDNSAGVDCSDHEVNIKILIDSLVSAGTVKADERTQLLESMTDEVAQLVLADNEDQNDLMGTSRANA
Sc/564-1650 EFDRNGGKVNTDAIDNSAGVDTSDHEVNIKILLNALVTEGDLTLKQRNILLAQMTDEVGALVLRNNYAQNVALANAVSQS
Pa/534-1619 EFGLHGGANNTDFIDNAGGVDCSDHEVNIKILLNEVVQAGDMTEKQRNALLVKMTDAVGALVLGNNYKQTQALSLAQRRA
Nf/55-1132 EYARAGGRINTDALDNSAGVDCSDHEVNIKILLDGLVGAGRLSTAERNALLAEMAAEVCRLVLADNESQNQLMGTARARA

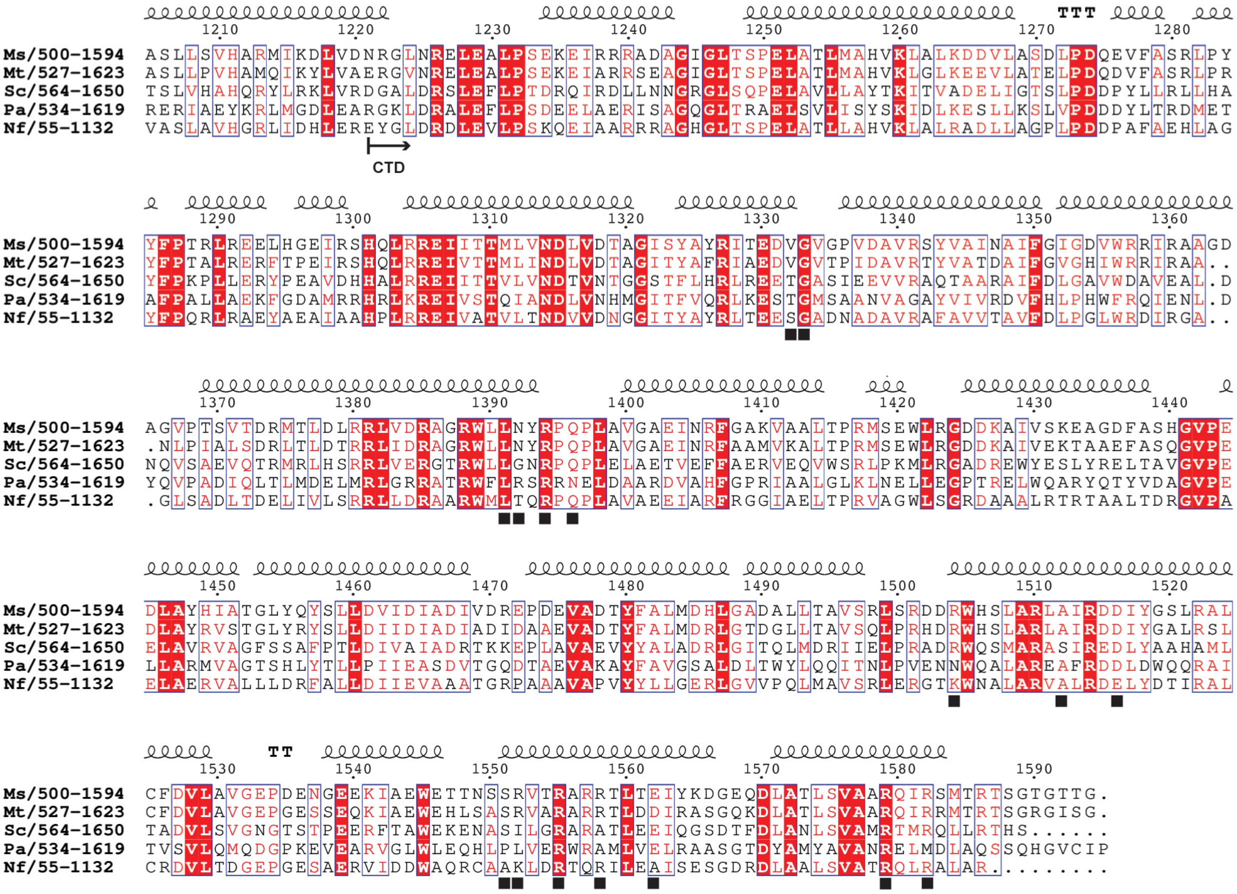


**Sequence alignment of diverse L-GDHs.** Sequences included: Ms, L-GDH$_{180}$ from *M. smegmatis* (UniProt A0R1C2); Mt, L-GDH$_{180}$ from *M. tuberculosis* (UniProt O53203); Sc, L-GDH$_{118}$ from *S. clavuligerus* (UniProt E2Q5C0); Pa, L-GDH$_{118}$ from *P. aeruginosa* (UniProt Q9HZE0) and L-GDH$_{115}$ *N. farcinica* (UniProt A0A0H5NTF9). The alignment was restricted to the region between residue 500 and the C-terminus of L-GDH$_{180}$ from *M. smegmatis*. Secondary structure elements derived from the atomic coordinates of the mL-GDH$_{180}$ monomer refined at 2.63 Å are shown above the alignment. The beginning of the domains of L-GDH$_{180}$ from *M. smegmatis*, as previously defined [1], are indicated with arrows. Residues involved in intersubunit salt bridges and/or hydrogen bonds in L-GDH$_{180}$ from *M. smegmatis* are marked with a square; active site residues are signaled with a circle. This figure was prepared using ESPript 3 (https://espript.ibcp.fr) [2].

**Figure S6**

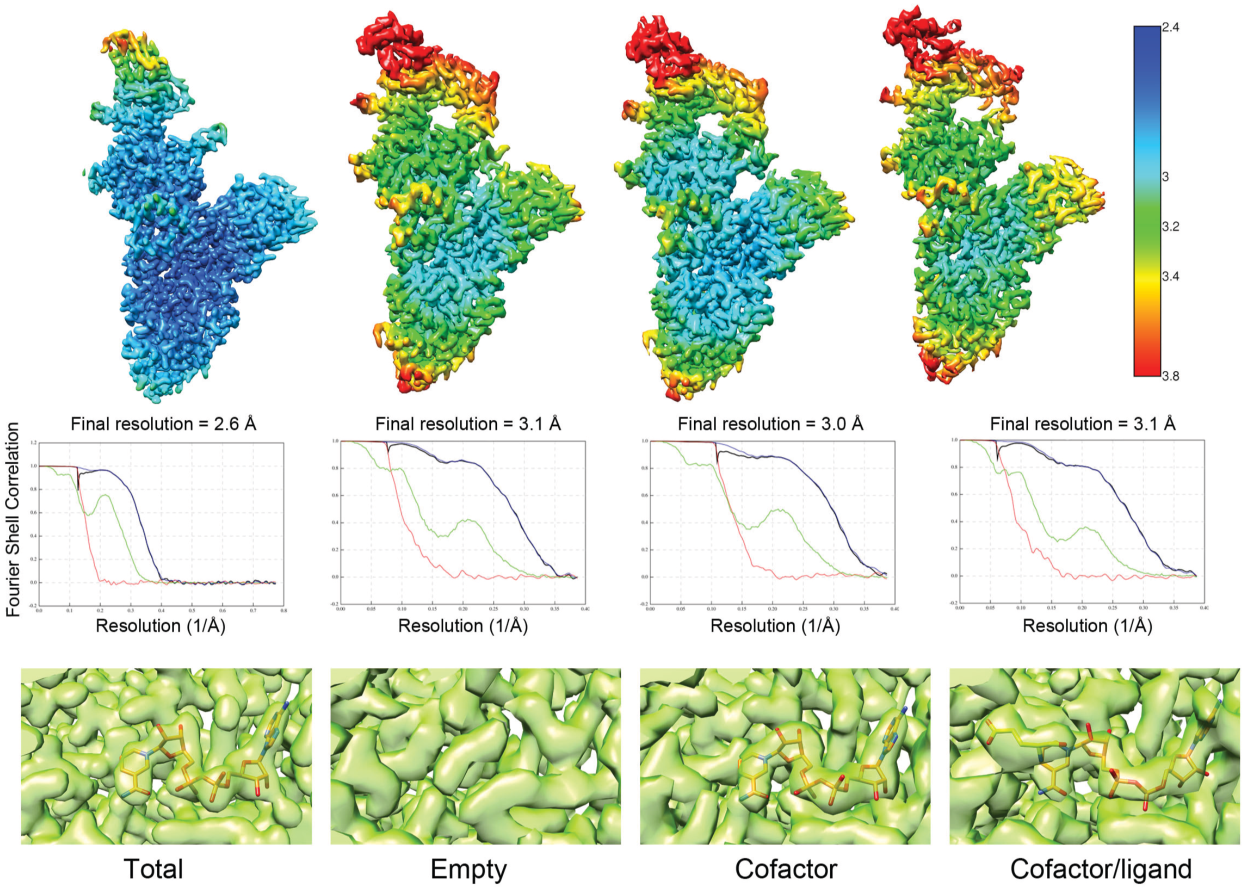


**Cryo-EM maps for mL-GDH$_{180}$ subunits.** Top panel: renderings for subunits colored by the estimated local resolution at the scale showed in the right side. Middle panel: Fourier Shell Correlation (FSC) graphs for each of the data sets. Lines are: red, FSC for phase randomized masked maps; green, FSC for unmasked maps; blue, FSC for masked maps; black, FSC for corrected and masked maps. Bottom panels: zoom-in views of the cofactor/ligand binding sites within the cryo-EM density maps for mL-GDH$_{180}$ subunits. The nature of the cryo-EM maps (total, empty, cofactor, and cofactor/ligand, see text) is labeled at the bottom of the figure.

**Figure S7**

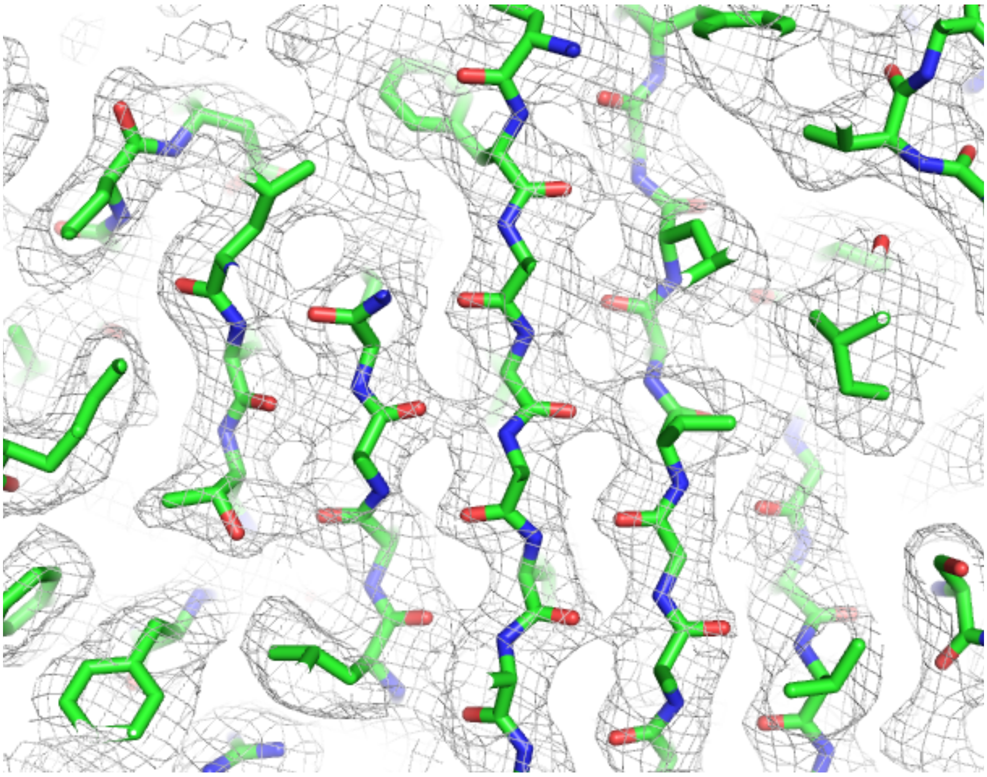

**A monomer of mL-GDH$_{180}$ at 2.63 Å.** The cryo-EM map is shown as a mesh (contour level = 0.005) and the atomic model is depicted in stick representation. The level of structural detail achieved can be appreciated.

#### SUPPLEMENTARY TABLES

**Table SI. Hydrogen bonds and salt bridges at the AB-CD and AD-BC interfaces in the different mL-GDH$_{180}$ conformers found in the presence of L-glutamate and NAD$^{+}$.**

| Open | Closed1 | Closed2 |
|---|---|---|
| **AB-CD interface** | | |
| **Hydrogen bonds** | | |
| Ser541 [sc]:[mc] Val556’ | Ser541 [sc]:[mc] Val556’ | Ser541 [sc]:[mc] Val556’ |
| Leu542 [mc]:[mc] Val556’ | Leu542 [mc]:[mc] Val556’ | Leu542 [mc]:[mc] Val556’ |
| | Ser543 [mc]:[sc] Gln550’ | |
| | Ser543 [sc]:[mc] Val556’ | Ser543 [sc]:[mc] Val556’ |
| | Gln550 [sc]:[mc] Ser543’ | |
| Val556 [mc]:[sc] Ser541’ | Val556 [mc]:[sc] Ser541’ | Val556 [mc]:[sc] Ser541’ |
| Val556 [mc]:[mc] Leu542’ | Val556 [mc]:[mc] Leu542’ | Val556 [mc]:[mc] Leu542’ |
| | Val556 [mc]:[sc] Ser543’ | Val556 [mc]:[sc] Ser543’ |
| Arg560 [sc]:[mc] Pro561’ | | |
| Pro561 [mc]:[sc] Arg560’ | | |
| | | Ser650 [sc]:[sc] Glu790’ |
| | | Glu790 [sc]:[sc] Ser650’ |
| **AD-BC interface** | | |
| **Salt bridges** | | |
| | Arg1504 [sc]:[sc] Glu1562’ | Arg1504 [sc]:[sc] Glu1562’ |
| Asp1516 [sc]:[sc] Arg1582’ | | Asp1516 [sc]:[sc] Arg1582’ |
| | Glu1562 [sc]:[sc] Arg1504’ | Glu1562 [sc]:[sc] Arg1504’ |
| Arg1582 [sc]:[sc] Asp1516’ | | Arg1582 [sc]:[sc] Asp1516’ |
| **Hydrogen bonds** | | |
| Val1332 [mc]:[sc] Arg1555’ | Val1332 [mc]:[sc] Arg1555’ | Val1332 [mc]:[sc] Arg1555’ |
| | Gly1333 [mc]:[sc] | |
| | | Gly1333 [mc]:[sc] |
| Leu1391 [mc]:[sc] | Leu1391 [mc]:[sc] | Leu1391 [mc]:[sc] |
| Asn1392 [mc]:[sc] | | Asn1392 [mc]:[sc] |
| | | Asn1392 [sc]:[sc] Arg1555’ |
| | Arg1394 [mc]:[sc] | Arg1394 [mc]:[sc] |
| Gln1396 [mc]:[sc] | | |
| Ala1512 [mc]:[sc] Arg1579’ | Ala1512 [mc]:[sc] Arg1579’ | Ala1512 [mc]:[sc] Arg1579’ |
| Ser1551 [sc]:[mc] | | |
| Arg1552 [sc]:[mc] | Arg1552 [sc]:[mc] | Arg1552 [sc]:[mc] |
| Arg1552 [sc]:[mc] | | Arg1552 [sc]:[mc] |
| | Arg1552 [sc]:[mc] | Arg1552 [sc]:[mc] |
| Arg1555 [sc]:[mc] Val1332’ | Arg1555 [sc]:[mc] Val1332’ | Arg1555 [sc]:[mc] Val1332’ |
| | Arg1555 [sc]:[mc] | |
| | | Arg1555 [sc]:[sc] Asn1392’ |
| | | Arg1558 [sc]:[mc] |
| Arg1579 [sc]:[mc] Ala1512’ | Arg1579 [sc]:[mc] Ala1512’ | Arg1579 [sc]:[mc] Ala1512’ |

sc, side chain; mc, main chain.
